# Integrating complementary biological information for multi-objective enzyme engineering

**DOI:** 10.64898/2026.09.15.751856

**Authors:** Nathaniel Blalock, Yvett Sosa, James Heuschkel, Ruihao Li, Xu Ma, Suttipol Radomkit, Hao Wu, Frederic Buono, Jeff Song, Noah Pefaur, Laura J Kingsley, Philip A Romero

**Affiliations:** Department of Chemical and Biological Engineering, University of Wisconsin-Madison, Madison, WI, USA; Department of Biomedical Engineering, Duke University, Durham, NC, USA; Biotherapeutics Discovery Department, Boehringer Ingelheim Pharmaceutical Inc., Ridgefield, CT 06877, USA; Drug Substance NCE US, Boehringer Ingelheim Pharmaceutical Inc., Ridgefield, CT 06877, USA

**Keywords:** multi-objective enzyme design, industrial biocatalysis, machine learning-guided, ketoreductase, protein engineering

## Abstract

Enzyme catalysts are increasingly used for sustainable pharmaceutical manufacturing, but engineering industrial biocatalysts remains challenging as multiple catalytic and developability properties must be optimized simultaneously from limited experimental data. Here, we develop a machine learning-guided multi-objective design framework that integrates sparse functional measurements with complementary evolutionary and structural information to engineer the ketoreductase Gre2. The resulting designs achieved simultaneous improvements in catalytic performance, protein yield, and thermal stability, with selected designs retaining improved performance under process-relevant conditions. More broadly, our results demonstrate that integrating complementary biological information enables efficient multi-objective enzyme engineering from sparse experimental data, providing a general strategy for accelerating industrial biocatalyst development.

## Introduction

Enzyme catalysts are important tools for the sustainable manufacture of pharmaceuticals and other high-value chemicals by enabling highly selective chemical transformations under mild reaction conditions^1–3^. Engineering enzymes to meet the stringent performance requirements for manufacturing processes remains a major protein engineering challenge^1,4^. Industrial applications require enzymes that simultaneously achieve high catalytic activity, selectivity, stability, expression, and robustness under process conditions^5,6^. Conventional directed evolution has achieved remarkable industrial successes, but the optimization of multiple enzyme properties is typically achieved through successive rounds of mutagenesis and screening, requiring substantial experimental effort and development time^7–9^. Strategies that simultaneously optimize multiple enzyme properties while reducing experimental iteration could substantially accelerate industrial biocatalyst development^2,10,11^.

Machine learning is increasingly used to guide protein engineering from experimental sequence-function data, offering a means to reduce screening requirements and accelerate protein engineering campaigns^12,13^. This is particularly valuable for industrial enzyme development, where process-relevant reactions are often difficult to implement at high throughput and development timelines permit only a limited number of sequence variants and experimental rounds. These sparse functional measurements sample only a small fraction of the sequence-function landscape, leaving most mutations and nearly all mutational combinations unobserved. Although supervised models can extrapolate beyond measured variants, prediction uncertainty increases as design moves farther from experimentally sampled sequence space^14,15^. Evolutionary and structural information can help constrain extrapolation into biologically plausible regions of sequence space. Evolutionary models trained on natural protein sequences capture constraints associated with natural enzyme function^16–18^, whereas structure-based models evaluate sequence compatibility with folded protein structures^19–22^. Integrating these complementary sources of information with sparse experimental data enables rapid prioritization of a small number of sequence designs that balance predicted activity, evolutionary plausibility, and structural compatibility, allowing experimental efforts to focus on a small number of candidates within limited development timelines.

Here, we develop an integrative framework for multi-objective enzyme design that combines three complementary models: a fine-tuned protein language model ESM-2 that learns reaction rate from sparse experimental measurements, a variational autoencoder trained on natural homologs that captures family-level evolutionary constraints, and SolubleMPNN that evaluates sequence compatibility with the enzyme structure while favoring soluble proteins. We apply this framework to the industrial ketoreductase, Gre2, capable of catalyzing the synthesis of a challenging chiral ketone intermediate through a kinetic resolution process^23,24^ (Supplementary Figure 1) by fine-tuning an ESM-2 model on 100 experimentally characterized variants and combining the resulting activity predictor with evolutionary likelihood and structural plausibility scores. Multi-objective optimization identified 1,219 variants predicted to improve all three computational objectives relative to Gre2, from which we selected 15 for experimental characterization. The resulting library was enriched for variants with simultaneous improvements in catalytic conversion, enantiomeric excess, protein yield, and thermal stability, while selected designs retained improved performance under process-relevant biocatalytic conditions. More broadly, our results demonstrate how complementary biological information can accelerate multi-objective protein engineering from sparse experimental data.

## Results

### Sparse mutational sampling identifies F85L as an activity-enhancing Gre2 variant

Industrial enzyme engineering campaigns often begin with experimental datasets that sparsely sample the sequence–function landscape. To establish an experimental foundation for low-data Gre2 engineering, we generated a single-mutant library using complementary knowledge-based, physics-based and protein language model approaches. Mutations were drawn from published mechanistic and structural evidence^25^, sequence alignments, aggregation-prone surface regions, ensemble-based residue scanning of apo- and holo-Gre2 structure conformations and substitutions ranked as evolutionarily plausible by a pre-trained protein language model^26^. Variants were selected to distribute sampling across the enzyme while enriching for substitutions predicted to preserve function or improve stability, thereby producing an informative dataset for subsequent engineering. The resulting dataset contained 99 Gre2 single mutants covering 67 of 342 sequence positions and 99 of 6,498 possible single-amino-acid substitutions. Thus, the dataset sampled 19.59% of Gre2 positions and only 1.52% of the full single-mutant sequence space (Supplementary Figure 2). Mapping the sampled positions onto the Gre2 structure showed broad but incomplete coverage across the enzyme scaffold (Figure 1a).

**Figure 1:**
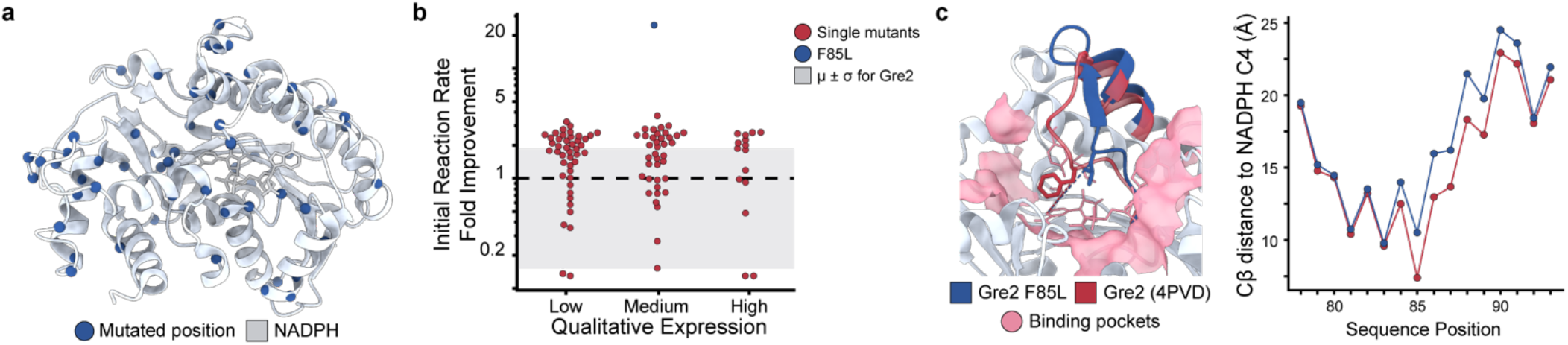
Sparse mutational sampling identifies F85L as an activity-enhancing Gre2 variant. **a**, Structural distribution of the initial Gre2 single-mutant training set mapped onto the Gre2–NADPH crystal structure^25^ (PDB: 4PVD). Sampled mutation positions are shown as blue spheres, with NADPH shown in gray. **b**, Initial reaction-rate measurements for the Gre2 single-mutant library, grouped by qualitative expression. The dashed line indicates Gre2 activity. **c**, Structural comparison of Gre2–NADPH^25^ (PDB: 4PVD) indicated in red and a predicted Gre2 F85L–NADPH model^27^ indicated in blue with the substrate and NADPH binding pockets indicated in pink. Alignment showed close global agreement, while F85L shifted the residue-85 Cβ atom farther from the NADPH nicotinamide C4 hydride-donor reference point, consistent with local active-site remodeling.

We characterized the 99 Gre2 single mutants and Gre2 for initial reaction rate and qualitative expression. F85L emerged as a beneficial mutation for reaction rate (Figure 1b; Supplementary Figure 3). This position was mechanistically plausible because prior structural work identified F85 as part of the hydrophobic pocket surrounding the substrate-binding channel, and NADPH binding was proposed to induce movement of the P84– C86 loop containing F85^25^. This suggested that a mutation at residue 85 could alter the geometry of the substrate-binding pocket. Consistent with this interpretation, structural comparison of Gre2–NADPH (PDB: 4PVD)^25^ with a predicted Gre2 F85L–NADPH model^27^ showed close global agreement after alignment, with a C*α* RMSD of 0.489 Å (Figure 1c; Supplementary Figure 4). Despite this close global agreement, the residue-85 Cβ atom in the predicted F85L model was positioned farther from the NADPH nicotinamide C4 atom, the hydride-donor carbon used as a mechanistic reference point. The Cβ-to-C4 distance increased from 7.39 Å in Gre2 to 10.51 Å in F85L, corresponding to a 3.12 Å increase. These results support a model in which F85L locally remodels the active site while preserving the overall Gre2 structure. We selected F85L as the parent sequence for subsequent multi-objective optimization.

### Benchmarking guides activity prediction from sparse mutational data

The sparse Gre2 dataset presented a practical modeling challenge: a supervised model needed to learn from limited measurements while still ranking untested variants for subsequent engineering. Developing an ESM-2– based predictor required choosing both how to represent each protein sequence for regression and how extensively to adapt the pretrained model to the experimental task. We therefore compared commonly used fixed-length sequence representations, including CLS-token and mean-pooled embeddings^13^, with position-specific full residue-level embeddings. We also evaluated fine-tuning strategies spanning linear regression on frozen ESM-2 embeddings^28^, LoRA-based parameter efficient fine-tuning with an MLP regression head^13,29^, and partial fine-tuning with an MLP regression head^30–32^.

Because the 99-variant Gre2 dataset was too small to support systematic model comparison, we benchmarked these representation and fine-tuning choices using external deep mutational scanning (DMS) datasets from CreiLOV^33^, avGFP^34^, and Ube4b^35^. For each protein, we constructed training sets containing single, double, triple, quadruple, or quintuple mutants, as well as a combined set spanning all five mutation orders, and evaluated performance on protein-specific held-out test sets (Supplementary Tables 1-3). This design allowed us to assess how representation and fine-tuning choices performed across variants with increasing numbers of mutations.

We first compared three representations derived from the final ESM-2 hidden layer (Supplementary Table 4). The CLS-token representation uses the embedding of a single sequence-summary token, whereas mean-pooling averages embeddings across the sequence, both producing a fixed-length representation. In contrast, the residue-level representation retains an embedding for every sequence position, preserving position-specific information at the cost of greater input dimensionality and computational expense. Across the benchmark datasets and held-out mutation regimes, residue-level representations generally achieved stronger predictive performance, particularly when models were trained on single-mutant data (Figure 2a; Supplementary Figure 5). These results suggest that retaining position-specific information is particularly useful in the sparse mutational regimes commonly encountered at the outset of industrial enzyme-engineering campaigns.

**Figure 2:**
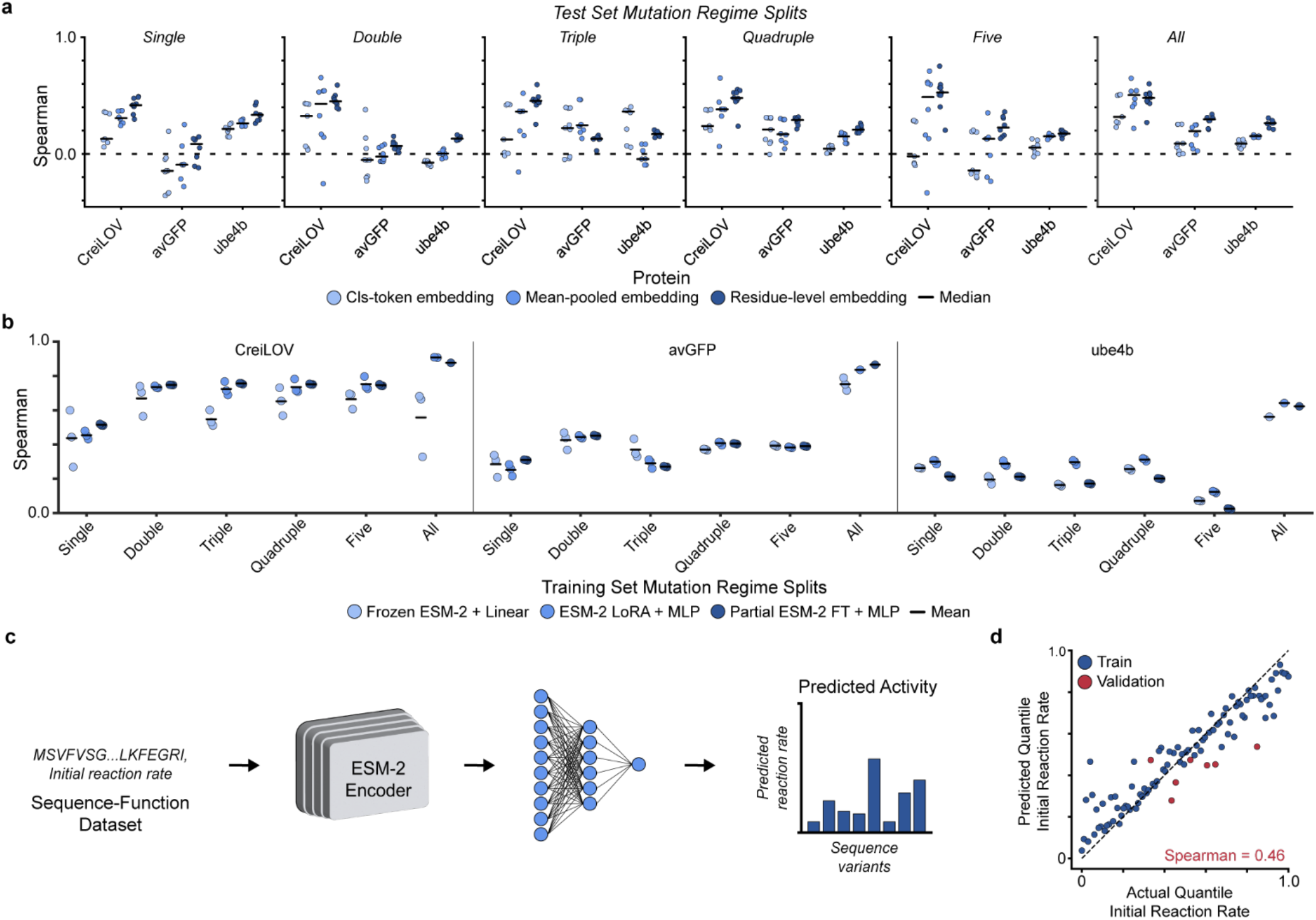
Benchmarking guides activity prediction from sparse mutational data. **a**, Benchmark of ESM-2 representation strategies across external DMS datasets. CLS-token, mean-pooled, and residue-level embeddings were compared across mutation-order training regimes, with residue-level embeddings generally improving low-data prediction. **b**, Benchmark of ESM-2 training strategies across the same external DMS datasets. Linear regression on frozen ESM-2 embeddings was compared with LoRA-based parameter-efficient fine-tuning and partial ESM-2 fine-tuning. Parameter-efficient fine-tuning frequently improved held-out prediction relative to frozen-embedding linear models. **c**, Supervised Gre2 activity-modeling workflow. ESM-2 was fine-tuned using the 99 characterized Gre2 single mutants to predict initial reaction rate from sequence. **d**, Validation performance of the fine-tuned Gre2 ESM-2 activity model. Validation set was utilized to define early stopping criteria during fine-tuning.

We next compared three training strategies that differed in the extent of task-specific model adaptation (Supplementary Table 5). The lowest-compute baseline used precomputed ESM-2 embeddings as inputs to a linear regression model without updating the language model. LoRA-based fine-tuning kept the ESM-2 backbone frozen while training low-rank adapters within selected attention layers together with an MLP regression head. Partial fine-tuning instead directly optimized a subset of pretrained ESM-2 parameters together with an MLP regression head. Across the external benchmarks, LoRA and partial fine-tuning frequently outperformed linear regression on frozen embeddings, indicating that limited task-specific adaptation of ESM-2 can improve sequence–function prediction from small, sparse training datasets (Figure 2b; Supplementary Figure 6).

Guided by these benchmarks, we selected a residue-level representation with partial ESM-2 fine-tuning and an MLP regression head to predict initial reaction rate from the 99 characterized Gre2 single mutants (Figure 2c; Supplementary Figure 7). The fine-tuned ESM-2 model achieved a validation Spearman correlation of 0.46 (Figure 2d), improving over the zero-shot ESM-2 baseline Spearman correlation of 0.16 (Supplementary Figure 8). We used the predicted reaction rate from this fine-tuned model as activity objective for downstream multi-objective sequence design.

### Integrating complementary models for multi-objective enzyme design

The supervised activity model provided an optimization objective for improving Gre2 reaction rate, but sequence optimization required extrapolating beyond the sparse experimental measurements used for training into a largely unexplored multi-mutant sequence space. In this regime, predictions become increasingly uncertain because the effects of most mutations and all mutational combinations have not been experimentally observed. Rather than relying on the finetuned ESM-2 supervised predictions alone, we integrated complementary evolutionary and structural information that is independent of the experimental activity measurements to guide exploration of sequence space.

We constructed a three-objective design framework that combined the finetuned ESM-2 supervised activity prediction with complementary evolutionary and structural information (Figure 3a-b). The supervised model predicts the engineering objective, Gre2 reaction rate, directly from sequence. Evolutionary information was obtained from a variational autoencoder trained on natural Gre2 homologs and provides a prior favoring variants consistent with family-level sequence constraints^15,17,36^. Structural information was obtained using SolubleMPNN^20,37^, which evaluates the compatibility of candidate sequences with the Gre2 fold while favoring soluble protein sequences. Together, these objectives integrate complementary functional, evolutionary, and structural information to guide multi-objective enzyme design.

**Figure 3:**
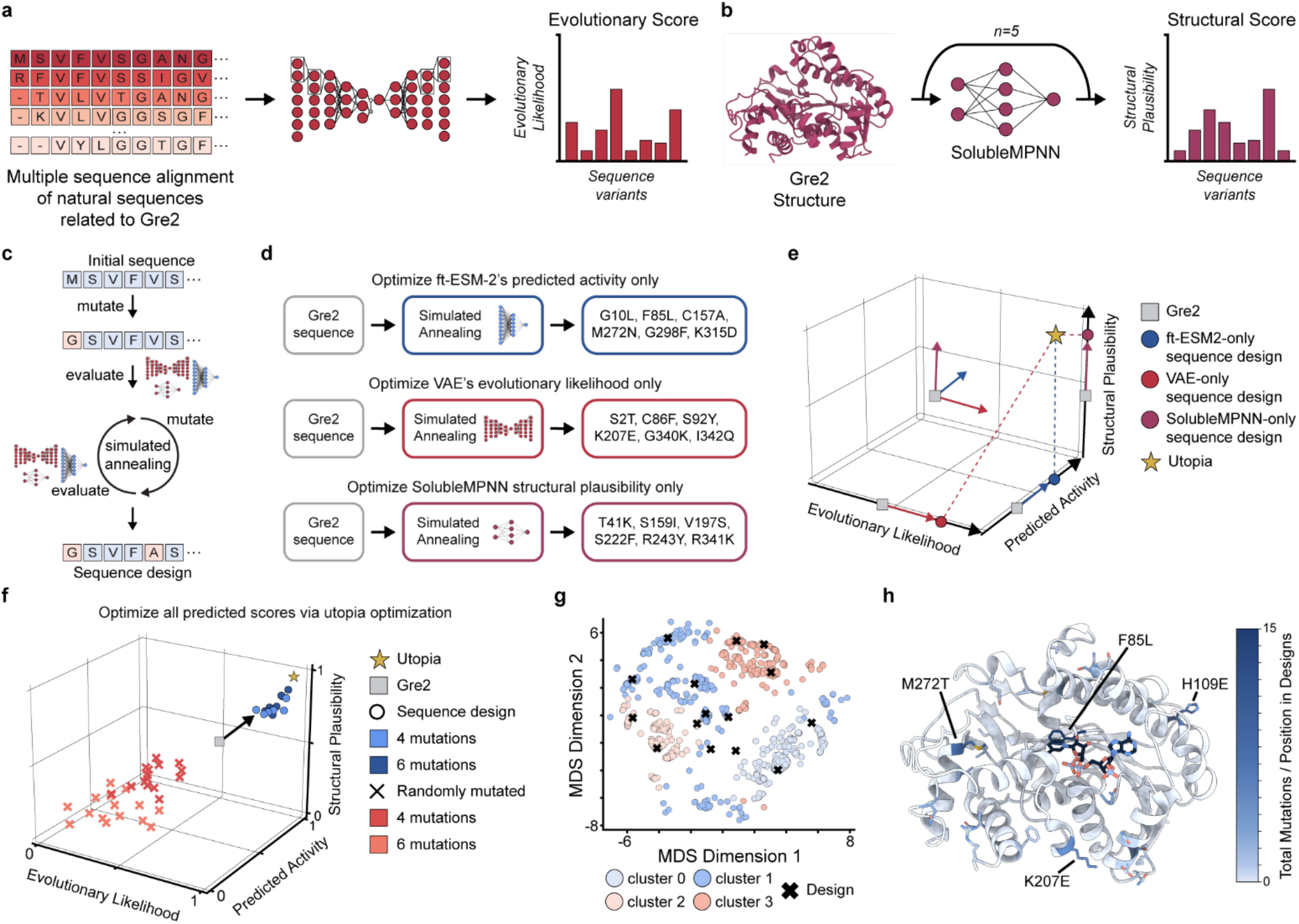
Integrating complementary models for multi-objective enzyme design. **a**, Variational autoencoder workflow for sequence-based scoring. A VAE was trained on a multiple sequence alignment of natural homologs related to Gre2 and used to predict the evolutionary likelihood of Gre2 sequence variants. This objective favors sequences that remain compatible with family-level constraints learned from natural enzyme diversity, providing an evolutionary prior complementary to supervised activity prediction. **b**, SolubleMPNN workflow for structure-based sequence scoring. Gre2 sequence variants were scored against the Gre2 structure to estimate compatibility. This objective provides a structural plausibility prior that is complementary to both the supervised activity predictor and the VAE evolutionary model. **c**, Simulated annealing strategy used to search Gre2 sequence space. Candidate sequences were iteratively mutated, evaluated by model scores, and accepted or rejected during optimization. **d**, Single-objective simulated annealing for each model objective. Fine-tuned ESM-2, the VAE, and SolubleMPNN were used to optimize the Gre2 sequence independently, producing distinct six-mutation Gre2 designs with no shared substitutions. **e**, Three-dimensional objective space showing the independently optimized model-specific designs and the theoretical utopia point defined by the best value for each objective. **f**, Multi-objective design by utopia optimization. Candidate variants were scored by their distance to the utopia point across predicted reaction rate, evolutionary likelihood, and structural plausibility. Randomly mutated variants are shown for comparison. **g**, Diversity-aware selection of multi-objective designs. High-scoring candidate sequences were clustered via K-means and sampled to avoid selecting near-duplicate variants. Final selected designs are shown as black X’s. **h**, Structural distribution of mutations selected in the final design library. Mutation frequency is mapped onto the Gre2 structure, with darker blue indicating positions mutated more frequently across the 15 selected designs. Frequently selected mutations included F85L, H109E, K207E, and M272T.

The evolutionary model distinguished natural Gre2-family sequences from random and heavily perturbed sequences, supporting its use as an evolutionary prior during sequence design (Figure 3a; Supplementary Figures 9-10). Likewise, introducing random mutations into Gre2 progressively reduced structural compatibility scores, and experimentally characterized variants with high qualitative expression tended to receive higher structural scores than random single mutants (Figure 3b; Supplementary Figure 11). These results support the use of evolutionary and structural objectives as complementary biological constraints that are independent of the experimental activity measurements.

To compare the design behavior of the three objectives, we separately optimized the Gre2 sequence using each model utilizing simulated annealing (Figure 3c,d). The optimized variants selected by each model had no shared mutations, demonstrating that activity prediction, evolutionary likelihood, and structural compatibility favor distinct regions of Gre2 sequence space (Figure 3d; Supplementary Figure 12). The finetuned ESM-2 activity model-optimized variant contained G10L, F85L, C157A, M272N, G298F, and K315D. The VAE-optimized variant contained S2T, C86F, S92Y, K207E, G340K, and I342Q. The SolubleMPNN-optimized variant contained T41K, S159I, V197S, S222F, R243Y, and R341K. Notably, only the activity-informed model selected F85L, consistent with F85L’s experimentally measured activity benefit. The VAE selected C86F in the adjacent P84–C86 loop but did not select F85L. This distinction is informative given that a predicted structure of the Gre2 homodimer^27^ placed Cys86 residues from opposing subunits in a geometry consistent with a potential interchain disulfide bond (Supplementary Figure 13). Gre2 homologs did not retain cysteine at this position (PDB: 7XPM, 5B6K^38^, and 7C3V), suggesting the VAE preference for C86F reflects broader family-level sequence variation rather than a Gre2-specific structural constraint as C86F would remove a potentially stabilizing interaction for the homodimer (Supplementary Figure 14). SolubleMPNN preferentially selected mutations that were more surface exposed (Supplementary Figure 12). This behavior is consistent with the idea that SolubleMPNN emphasizes fold compatibility and soluble-protein-like surface composition rather than catalytic activity.

Because each model captures a different aspect of enzyme fitness, no single sequence is expected to simultaneously maximize all three objectives. We therefore formulated sequence design as a multi-objective optimization problem. First, each objective was optimized independently to identify its theoretical optimum for predicted activity, evolutionary likelihood, and structural compatibility (Figure 3c-d). These independent optima define a “utopia” point representing the best possible combination of all three objectives in the absence of tradeoffs (Figure 3e). Candidate sequences were then prioritized according to their Euclidean distance from this utopia point, favoring designs predicted to improve activity while remaining evolutionarily plausible and structurally compatible.

We then used simulated annealing to search Gre2 sequence space under the utopia-distance objective (Figure 3f). Designs were generated from the F85L parent, thereby retaining the experimentally validated activity-enhancing mutation while exploring additional substitutions predicted to improve the combined activity, evolutionary, and structural objectives. This search identified 1,219 sequence variants with all three computational scores improved relative to Gre2. Because direct optimization can converge on closely related local optima, we clustered high-scoring sequences and applied cluster-aware sampling to select a diverse experimental set rather than near-duplicate variants (Figure 3g). This procedure yielded a final library of 15 multi-mutant Gre2 designs for experimental characterization. The most frequently added mutations in the context of F85L included H109E, K207E, and M272T with additional mutations sampled across the enzyme structure (Figure 3h).

### Machine learning-guided design improves multiple industrially relevant properties

To test whether computational improvements translated into experimental performance, we synthesized the 15 diverse Gre2 variants predicted to improve all three model objectives and experimentally characterized these variants for four process-relevant properties: conversion, enantiomeric excess, protein yield measured as total protein produced from the expression culture, and thermal stability measured by melting temperature (Tm) (Figure 4a-d, Supplementary Table 6). The reaction is a kinetic resolution of a 1:1 mixture of R- and S-ketones in which the KRED preferentially reduces the S-ketone to the S-alcohol while the desired R-ketone remains unreacted and is recovered (Supplementary Figure 1). For a perfectly selective reaction, the theoretical maximum conversion is therefore 50%. Because enantiomeric excess was measured for the residual ketone substrate, it is inherently correlated with conversion: as the S-ketone is consumed, the unreacted R-ketone is enriched in the substrate mixture. Accordingly, a higher residual-ketone enantiomeric excess value may reflect greater conversion and does not directly demonstrate improved intrinsic enantioselectivity. We therefore interpret residual ketone enantiomeric excess as enrichment of the recovered R-ketone before the reaction reaches 50% conversion. Together, these measurements capture both reaction performance and developability, allowing us to evaluate whether the computational design strategy improved catalytic properties without sacrificing expression or biophysical robustness.

**Figure 4:**
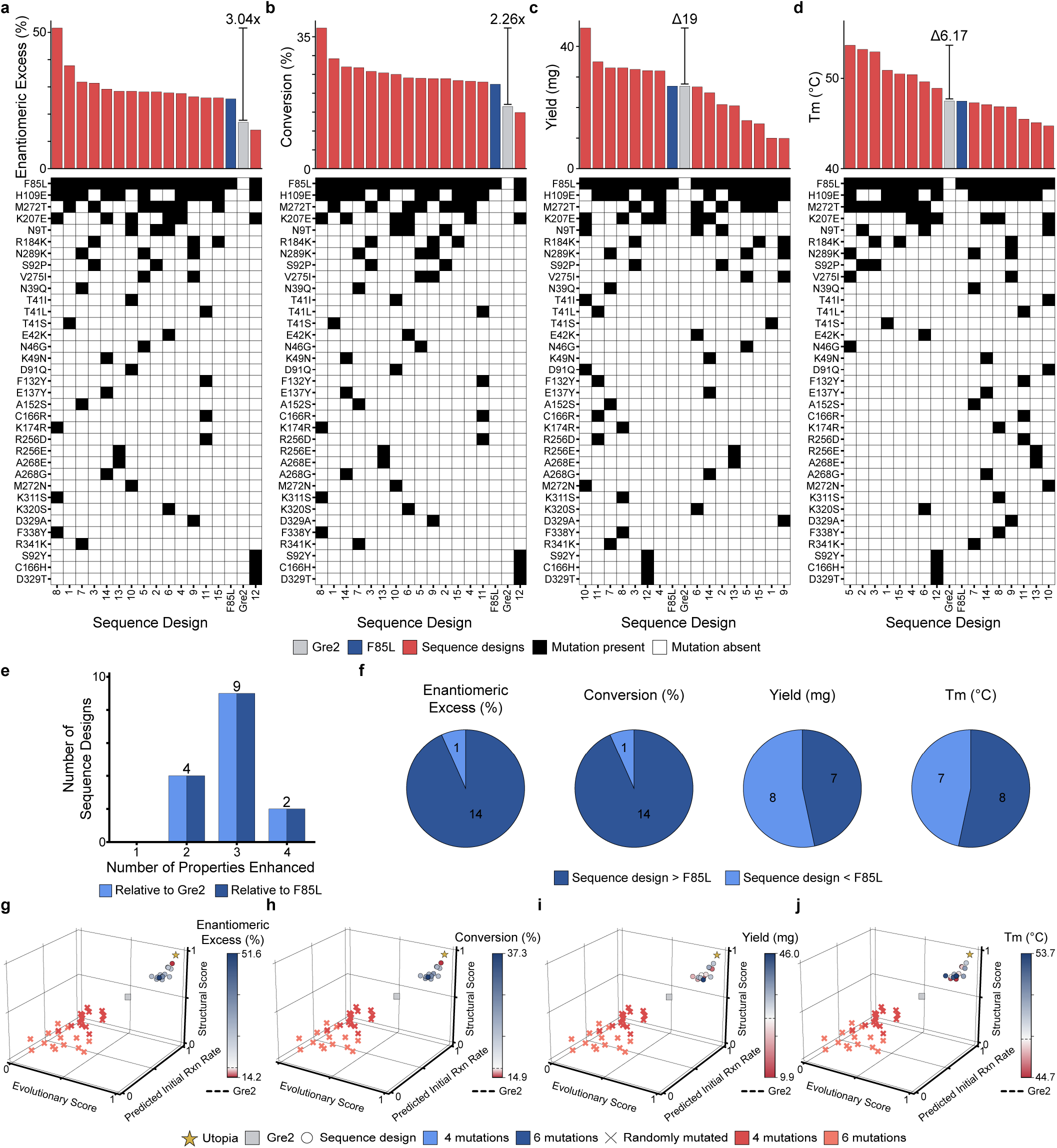
Machine learning-guided design improves multiple industrially relevant properties. **a-d**, Experimental characterization of 15 multi-objective Gre2 sequence designs for four process-relevant properties: enantiomeric excess, conversion, protein yield, and thermal stability. Gre2, F85L, and sequence designs are shown for comparison. **e**, Number of measured properties improved by each design relative to Gre2 and the F85L parent. **f**, Fraction of designs improved relative to F85L for each property. **g-h**, Ranked enantiomeric excess and conversion values with corresponding mutation heatmaps. Sequence design 8, containing F85L, H109E, K174R, K207E, K311S, and F338Y, showed the strongest reaction performance, reaching 51.6% enantiomeric excess and 37.3% conversion. **i**, Ranked protein yield with mutation heatmap. Sequence design 10, containing N9T, T41I, F85L, D91Q, K207E, and M272N, produced the highest yield of 46 mg/L. **j**, Ranked melting temperature with mutation heatmap. Sequence design 5, containing N46G, F85L, H109E, M272T, V275I, and N289K, showed the highest thermal stability, reaching a Tm of 53.7 °C. Mutation heatmaps indicate the substitutions present in each design, highlighting that the best variants for reaction performance, yield, and thermostability used distinct mutation combinations beyond the shared F85L parent mutation.

Eleven of 15 designs improved at least three of four measured properties relative to both Gre2 and the F85L parent sequence (Figure 4e). Improvements in reaction performance were especially common: 14 of 15 designs improved both conversion and enantiomeric excess relative to F85L (Figure 4f). Expression yield and thermal stability improved in a substantial fraction of the library, with 7 of 15 designs improving yield and 8 of 15 designs improving melting temperature relative to F85L (Figure 4f). Thus, the multi-objective strategy achieved simultaneous improvements across reaction performance and developability.

The strongest variant for both conversion and enantiomeric excess was sequence design 8, containing F85L, H109E, K174R, K207E, K311S, and F338Y (Supplementary Figure 15-16). This variant reached 51.6% enantiomeric excess and 37.3% conversion, corresponding to 3.0-fold and 2.3-fold improvements relative to Gre2 (Figure 4a-d; Supplementary Figure 17). The variant with the best yield was sequence design 10, containing N9T, T41I, F85L, D91Q, K207E, and M272N (Supplementary Figure 15-16). This design produced 46 mg/L of protein, a 19 mg/L increase relative to Gre2 (Figure 4c; Supplementary Figure 17). The variant with the best thermostability was sequence design 5, containing N46G, F85L, H109E, M272T, V275I, and N289K (Supplementary Figure 15-16). This variant reached a melting temperature of 53.7°C, corresponding to a 6.17°C increase relative to Gre2 (Figure 4d; Supplementary Figure 17). Notably, the gains across these properties, including 3.0-fold higher enantiomeric excess value and 2.3-fold higher conversion, were identified by testing only 15 sequence designs in a single design-test cycle, highlighting the potential to reduce experimental screening and development time.

Although the design library was enriched for improved variants, experimental performance did not map perfectly onto any single computational score or onto Euclidean distance to the utopia point (Supplementary Figure 18). This is not unexpected in a highly optimized multi-objective region of sequence space, where many candidate designs are predicted to satisfy the computational filters but exhibit different property tradeoffs and background-dependent mutational effects from epistasis. This result supports the use of diversity-aware sampling after multi-objective optimization rather than selecting only the nearest neighbors to a single computational optimum (Figure 3g, 4g-j).

We next examined mutational patterns among the best-performing variants to assess how multi-objective design produced improved enzymes. The top variants did not converge on a single shared mutational solution beyond the F85L parent mutation. Instead, variants with the best reaction performance, yield, thermostability, or balance across all four properties contained distinct additional mutation sets (Figure 4a-d, Supplementary Figure 15-16).This suggests that Gre2 improvement can be achieved through multiple sequence routes and that the different measured properties do not share a single dominant mutational optimum.

Several recurrent positions nevertheless emerged across high-performing designs. H109E was present in the variants with the best reaction performance and thermostability and the variant with the second-best yield (Figure 4a-d). K207E was present in the variants with the best reaction performance and yield and the variant with the sixth-best thermostability (Figure 4a-d). Both H109E and K207E are positioned on helices surrounding the NADPH-binding pocket (Supplementary Figure 15-16). K207E is also located near the modeled homodimer interface that helps form the mouth to the NADPH-binding pocket (Supplementary Figure 19). Because NADPH binding has been proposed to promote interdomain motion and formation of a substrate-binding cleft, these cofactor-proximal substitutions may influence local packing, cofactor-associated conformational changes, or interdomain interactions^25^.

Position M272 showed a substitution-dependent relationship with yield and thermostability. M272N was present in sequence design 10, the variant with the best yield, whereas M272T was present in sequence design 5, the variant with the best thermostability (Figure 4c-d; Supplementary Figure 15-16). This suggests that substitutions at M272 may tune multiple properties, but that different amino acids favor different experimental outcomes. Consistent with this interpretation, design 10 had the highest yield but the lowest Tm, suggesting a possible yield–thermostability tradeoff associated with its mutation set, potentially involving T41I, D91Q, and/or M272N. By contrast, the highest Tm was achieved by a distinct combination of N46G, H109E, M272T, V275I, and N289K. These patterns support a background-dependent model in which property optima occupy different regions of Gre2 sequence space rather than arising from a universally beneficial mutation set.

Sequence design 3 provided a useful example of balanced multi-objective improvement. This variant contained only four mutations, F85L, S92P, R184K, and M272T, and improved all four measured properties without being the top design for any single metric. S92P is positioned near the F85-containing active-site region, M272T lies near the substrate-binding region, and R184K is a conservative charge-preserving substitution (Supplementary Figure 15-16). The presence of M272T in both design 3 and the variant with the best thermostability suggests that M272T may be broadly compatible with improved developability and reaction performance. More generally, design 3 supports the premise that multi-objective design can identify compact mutation combinations that improve catalytic performance while preserving or improving biophysical properties.

Together, these results show that ML-guided design can produce variants with simultaneous improvements in catalytic performance and developability. The experimentally improved variants were not explained by a single mutation or structural region. Instead, the best reaction-performance, yield, thermostability, and balanced variants sampled distinct combinations of active-site-proximal, cofactor-proximal, substrate-proximal, and scaffold-distributed mutations. Recurrent positions such as H109, K207, and M272 point to structural regions that may be useful for balancing multiple properties, but the limited overlap among top variants suggests that Gre2 improvement is distributed and background dependent. This supports the central premise of the multi-objective strategy: combining activity prediction with evolutionary and structural priors can identify diverse mutation combinations that improve catalytic performance while preserving or improving developability.

### Designed enzymes retain performance under process-relevant conditions

Industrial biocatalytic reactions often require operating conditions that differ from standard screening assays. Optimizing reaction conditions, including pH and temperature, can improve substrate availability, reaction rate, or compatibility with the broader manufacturing process^7–9,39^, while elevated temperatures can reduce contamination risk in whole-cell systems^40^. However, these conditions can also reduce enzyme activity, selectivity, or operational stability^7–9,39^. We therefore asked whether the multi-objective sequence designs retained improved performance under pH and temperature challenges relevant to process development. Selected designs were assayed across varying pH and temperature conditions, and conversion and enantiomeric excess were measured as reaction-performance outputs (Figure 5).

**Figure 5:**
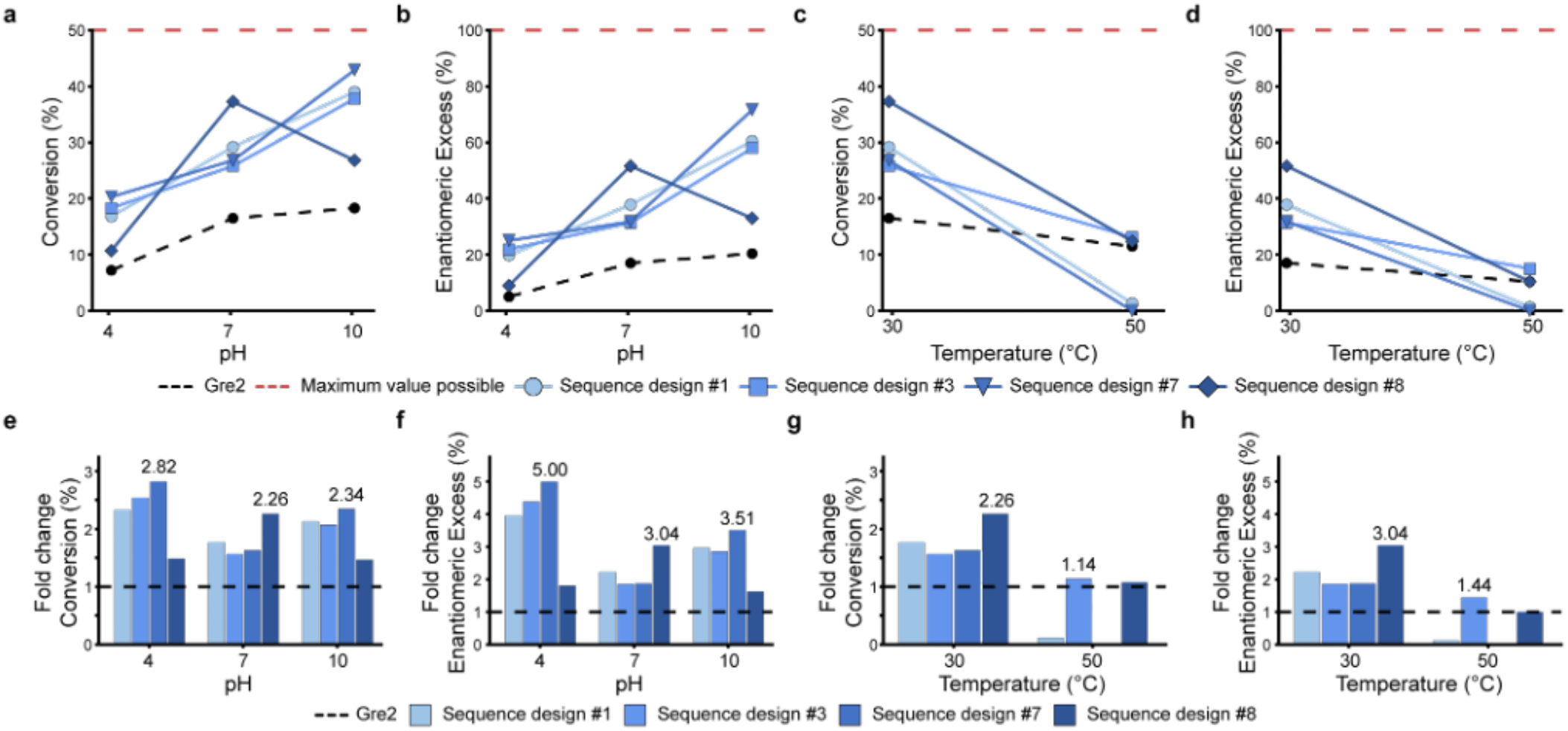
Designed enzymes retain performance under process-relevant conditions. **a**, Conversion of selected sequence designs across pH challenge conditions. **b**, Enantiomeric excess of selected sequence designs across pH challenge conditions. **c**, Conversion of selected sequence designs across temperature challenge conditions. **d**, Enantiomeric excess of selected sequence designs across temperature challenge conditions. Red dashed lines indicate the maximum possible values: 50% conversion in (a) and (c) and 100% enantiomeric excess in (b) and (d). Black dashed lines indicate Gre2 performance. **e**, Conversion fold change of selected sequence designs across pH challenge conditions. Dashed line indicates wildtype GRE2 performance. **f**, Enantiomeric excess fold change of selected sequence designs across pH challenge conditions. Dashed line indicates wildtype GRE2 performance. **g**, Conversion fold change of selected sequence designs across temperature challenge conditions. Dashed line indicates wildtype GRE2 performance. **h**, Enantiomeric excess fold change of selected sequence designs across temperature challenge conditions. Dashed line indicates GRE2 performance.

We first evaluated pH tolerance range. Selected multi-objective sequence designs were tested at pH 4, pH 7, and pH 10. Across all three pH conditions, the designs improved conversion and enantiomeric excess value relative to Gre2 (Figure 5a,b; Supplementary Table 7). The best variants showed conversion improvements of 2.8-fold, 2.3-fold, and 2.3-fold at pH 4, pH 7, and pH 10, respectively (Figure 5e). Enantiomeric excess value was also improved across the pH range, with best observed improvements of 5.0-fold, 3.0-fold, and 3.5-fold at pH 4, pH 7, and pH 10, respectively (Figure 5f). These results indicate that the improved reaction performance of the multi-objective sequence designs was not restricted to the original screening pH.

We then evaluated robustness to temperature. Selected multi-objective sequence designs were assayed at 30 °C and 50 °C. At 30 °C, all tested designs outperformed Gre2 for both conversion and enantiomeric excess (Figure 5c,d; Supplementary Table 8), with best observed improvements of 2.3-fold for conversion and 3.0-fold for enantiomeric excess value (Figure 5g,h). At 50 °C, overall performance decreased, consistent with the increased severity of the condition. Nevertheless, designs 3 and 8 retained improved performance relative to Gre2, with best observed improvements of 1.14-fold for conversion and 1.44-fold for enantiomeric excess value (Figure 5g,h). Thus, while high temperature remained challenging, a subset of sequence designs maintained reaction performance even under elevated-temperature conditions.

The strongest process-challenge result was observed for sequence design 7 under alkaline and moderate-temperature conditions. At pH 10 and 30 °C, design 7 reached 42.9% conversion and 71.6% enantiomeric excess, the highest combined reaction performance observed in the challenge assays (Figure 5). This result is notable because it shows that the multi-objective sequence designs can perform well outside the initial characterization condition and suggests that they provide improved starting points for further process optimization.

Together, these pH and temperature challenge experiments show that selected multi-objective sequence designs retain improved conversion and enantiomeric excess across process-relevant reaction environments. These experiments do not constitute full process development, but they demonstrate the designed variants are more robust starting points than Gre2 under several nonstandard assay conditions. This supports the broader premise that combining activity prediction with evolutionary and structural priors can produce enzyme variants with improved catalytic performance and greater process tolerance.

## Discussion

Sparse experimental measurements provide an incomplete description of enzyme fitness, limiting the ability of supervised models to guide sequence optimization far beyond observed variants. Here, we show that integrating complementary biological information with sparse functional measurements enables effective multi-objective enzyme engineering. Starting from only 99 characterized Gre2 single mutants and Gre2, the framework identified variants with simultaneous improvements in catalytic performance and developability. More broadly, these results demonstrate that complementary evolutionary and structural information can guide sequence optimization in regions of sequence space not represented in the experimental training data.

A central challenge in industrial biocatalysis is that useful enzymes must satisfy several constraints simultaneously. Improved reaction rate alone is not sufficient if the variant expresses poorly, loses solubility, unfolds, or fails under process-relevant pH, temperature, solvent, or substrate-loading conditions. This limitation was evident in the initial screen, where F85L improved reaction rate but did not improve qualitative expression. In contrast, 11 of 15 multi-objective designs improved at least 3 of 4 measured properties relative to both Gre2 and F85L, and 14 of 15 improved both conversion and enantiomeric excess relative to F85L. Protein yield and thermostability, which were not directly supervised during design, also improved in 7 of 15 and 8 of 15 designs, respectively. Thus, optimizing activity in the context of evolutionary and structural constraints enriched the library for catalytic improvements without sacrificing developability.

The concentration of improved variants within a small experimental set of 15 sequence designs highlights the efficiency of this approach. Across the full campaign, comprising initial training-set generation followed by one model-guided design–build–test cycle, we characterized 115 sequence identities: Gre2, 99 single mutants, and 15 multi-mutant designs. Among the 15 designs, we identified a variant containing N39Q, F85L, H109E, A152S, N289K, and R341K that reached 42.9% conversion and 71.6% enantiomeric excess at pH 10 and 30 °C, corresponding to 2.3-fold and 3.5-fold improvements over Gre2, respectively. For context, a sitagliptin transaminase campaign screened 36,840 variants across nine sequential optimization rounds to adapt the enzyme to process conditions, averaging approximately 4,093 variants per round^7^. An industrial ketoreductase campaign screened 15,025 transformants, representing an estimated 6,607 unique sequence variants, across 6 evolution rounds, averaging approximately 2,504 transformants or 1,101 unique on-target variants per round^9^. In comparison, the average sitagliptin round therefore screened approximately 273 times as many variants as the Gre2 model-guided round, whereas the average ketoreductase round evaluated approximately 73 times as many estimated unique on-target variants. Although differences in enzymes, substrates, assays, and optimization goals preclude direct comparisons of improvement, these studies contextualize the substantially lower wet-lab screening burden of the model-guided Gre2 round. Our results suggest that integrating complementary computational objectives may reduce screening requirements and shorten enzyme-development campaigns while preserving or improving protein yield and thermostability.

The modeling benchmarks provide practical guidance for learning from sparse datasets typical of industrial biocatalysis. Representation choice determines the level of sequence detail made directly available to the downstream predictor. Residue-level representations generally outperformed CLS-token and mean-pooled embeddings in our benchmarks, with the clearest advantage when training was restricted to single mutants. This advantage may arise because CLS-token and mean-pooled embeddings compress a protein into a fixed-length vector by routing information through one special token or averaging across all residues. Although computationally efficient, this compression can obscure localized mutational signals in sparsely sampled sequence–function datasets. Prior protein-language-model studies similarly indicate that representation performance is task dependent. Across 40 DMS datasets, mean pooling performed best on average among the tested compression strategies, including CLS-token and max-pooled embeddings^41^. Notably, residue-level representations were not evaluated. However, learned residue weighting outperformed simple pooling for protein localization tasks^42^. Similar context dependence has been observed in NLP. After BERT introduced the CLS token as an aggregate representation for task-specific fine-tuning^43^, it was later shown that CLS-token and mean-pooled BERT embeddings performed poorly for cosine-based semantic similarity but remained reasonably effective as inputs to a trained classifier^44^. Together, these findings suggest that representation choice should reflect the downstream task and data regime. In the sparse single-mutant regime examined here, retaining complete position-specific information was most effective. If full residue-level representations are computationally impractical, learned position-aware aggregation or projection may provide an efficient intermediate strategy.

The fine-tuning strategy determines how strongly pre-trained representations are adapted to the measured phenotype. Linear regression on frozen embeddings assumes that assay-relevant information is already linearly accessible in the final ESM-2 layer. LoRA learns constrained low-rank updates, whereas partial fine-tuning updates selected pretrained parameters without imposing a low-rank constraint. The frequent improvement of both strategies over frozen-embedding regression agrees with external systematic pLM benchmarks showing that supervised fine-tuning improves diverse protein predictions^13^. Together, these findings indicate that representations learned through masked-residue prediction can benefit from limited adaptation to an experimentally measured phenotype. These conclusions should nevertheless be interpreted as practical guidance rather than universal rules given the benchmarks covered only three proteins, one ESM-2 model size, and a limited set of representations and fine-tuning strategies.

Related approaches have incorporated evolutionary or structural information into protein optimization, although generally with different objectives. Evolutionary and structure-based regularization has been applied separately in Bayesian optimization to discourage improvement of a measured property at the expense of unmeasured properties^45^. MODIFY uses an ensemble of unsupervised evolutionary models to co-optimize zero-shot predicted fitness and library diversity that is followed by structure-based filtering^11^, whereas the present framework explicitly balances an experimentally supervised activity predictor with separate evolutionary and structural objectives. The utopia-based formulation is also extensible to additional objectives that can be scored and normalized, including direct predictors of expression, thermostability, solvent tolerance, substrate scope, stereoselectivity, or manufacturability. Combined with diversity-aware sampling, this provides a strategy for sampling diverse solutions predicted to improve all objectives rather than converging on a single computational optimum.

The mutational patterns among the experimentally improved variants further support a distributed model of Gre2 improvement. The best reaction-performance, yield, thermostability, and balanced all-properties-enhanced variants shared F85L but otherwise used distinct mutation combinations. Recurrent positions such as H109, K207, and M272 point to structural regions that may help balance multiple properties, but the limited overlap among top variants suggests that their effects are background dependent. This is consistent with the broader premise of multi-objective design that different properties can require different sequence features, and a useful design strategy should sample diverse solutions rather than collapse onto a single mutational route.

The pH and temperature experiments indicate that the designs were not improved only under the initial screening condition. Selected multi-objective sequence designs improved conversion and enantiomeric excess across pH 4, 7, and 10, and all tested designs outperformed Gre2 at 30 °C. At 50 °C, overall performance decreased, but 2 designs retained improved performance relative to Gre2. These assays do not constitute full process optimization, but they indicate that the designs provide improved starting points for further development. Several limitations remain. Although the best design reached 42.9% conversion, 85.8% of the 50% theoretical maximum imposed by the reaction scheme, further improvements in enantiomeric excess and robustness across process conditions would be required for a final process. The pH and temperature experiments test process-relevant stressors but do not cover the full range of industrial variables, such as enzyme loading, substrate concentration, or solvent tolerance. Structural interpretations are hypotheses based on modeling and mutation co-occurrence and will require mutational dissection and experimental structural analysis. Finally, the evolutionary and structural scores remain proxies for unmeasured aspects of enzyme fitness. Future rounds could use the multi-mutant data generated here to train direct predictors of conversion, enantiomeric excess, yield, and melting temperature and to evaluate whether iterative active learning produces further improvements.

Together, these results show that machine learning-guided multi-objective design can use sparse experimental data to generate variants with improved catalytic and process-relevant properties. By combining an assay-trained activity predictor with evolutionary and structural priors, the workflow identified diverse mutation combinations that improved Gre2 reaction performance while preserving or improving yield and thermostability. This strategy provides a practical route to reducing the experimental burden and development time required to generate improved starting points for industrial enzyme optimization.

## Methods

### Selecting Gre2 single mutants

The initial library of 99 single-point mutants was designed using a combination of knowledge-based, structure-based, and pLM approaches. The goal of the 100 mutant library was to obtain a first training data set made up of functional mutations that broadly span the whole protein sequence and have varying functional data. Knowledge-based mutations were based on sequence alignment and a seminal paper of GRE2^25^ which highlights key residues in the substrate-binding pocket that influence substrate stereoselectivity. Molecular dynamics simulations of the apo- (4PVD) and holo- (4PVC) Gre2 structures were performed using the Schrödinger Software suite. Each trajectory was grouped into three clusters and subsequently used for single-point mutational residue scanning using CCG software MOE (Molecular Operating Environment). Single point mutations at selected residues were ranked according to their predicted effects on protein stability (ΔΔGstability) and binding affinity (ΔΔGaffinity). We also evaluated the sequence using pLM based affinity maturation^26^.

### Production of Gre2 single mutant library

Expression constructs encoding Gre2 and Gre2 variant sequences were synthesized without secretion, detection, or solubility tags. The genes were cloned into a pET28-like vector under the control of an isopropyl β-D-1-thiogalactopyranoside-inducible promoter and expressed in an Escherichia coli BL21(DE3)-equivalent host. Following expression, cells were harvested and disrupted, and insoluble material was removed to obtain clarified cell lysates. The clarified lysates were lyophilized and stored at -20 ºC until use.

### Characterization of Gre2 single mutant library

Ketoreductase activity in lyophilized Escherichia coli cell lysates was measured by monitoring NADPH depletion at 340 nm. Lysates were reconstituted at 2.5 mg mL−1, and reactions were performed in 96-well black, clear-bottom plates (Corning, cat. no. 3880) in a final volume of 100 µL. Reactions contained 0.05 g/L lysate, 1.25 mM ketone substrate, 1.7 mM MgSO4, 1 mM NADPH, 122 mM sodium phosphate (pH 7.0), and 5% (v/v) dimethyl sulfoxide. Reactions were performed at pH 7.

Reactions were initiated by adding lysate, centrifuged at 500 rpm for 30 seconds, and mixed at 800 rpm for 30 seconds. Absorbance was recorded at 25 °C every minute for 2 h using a microplate reader with continuous slow double-orbital shaking. Wild-type ketoreductase and assay buffer or green fluorescent protein-expressing E. coli lysate served as positive and negative controls, respectively. Samples were measured in triplicate, and activity was calculated from the linear decrease in absorbance at 340 nm. Data was then analyzed as increase over parent fold improvement of reaction rates.

Ketoreductase expression in clarified cell lysates was assessed qualitatively by SDS–PAGE based on the presence and relative intensity of a protein band at the expected molecular weight.

### Production of Gre2 and Gre2 multi-objective designed mutants

The coding sequence for GRE2 and GRE2 mutants, respectively, was cloned into a pET28a-derived expression vector, adding an N-terminal hexahistidine-tag to the protein. The proteins were expressed in E. coli BL21(DE3) cells in 2×YT medium. The expression was performed in a bioreactor under controlled fermentation conditions to ensure a similar expression profile for all the protein variants (induction with 0.2 mM IPTG at an OD600 = 0.6 and recombinant protein expression for 20 hours at 16°C). Cells were harvested by centrifugation, and cell pellets were stored at −20°C. Cells were lysed in 20 mM Tris-HCl (pH 7.0), 200 mM NaCl and the clarified lysate was used for purification of GRE2 with immobilized metal affinity chromatography (elution with 20 mM Tris-HCl (pH 7.0), 200 mM NaCl, 500 mM imidazole) and a subsequent desalting column (20 mM Tris-HCl (pH 7.0), 200 mM NaCl). Fractions were pooled and concentrated. Protein expression and purity were evaluated by SDS-PAGE and Western blotting.

### Enzymatic Assays

NADPH depletion enzymatic assay reactions (100 µL) contained 10 µL E. coli lysate, 1.25 mM substrate, 1.7 mM MgSO_4_, 1 mM NADPH, and 122 mM Na_3_PO_4_. Plates (96-well clear-bottom) were incubated for 2 h and absorbance was measured at 340 nm. Small-scale biocatalytic reactions (200 µL) were performed in 1.5 mL Eppendorf tubes contained 50 mM substrate, 20 mM NADH or NADPH, and 5 µM purified enzyme in 100 mM NaPi (pH 7.0) with 150 mM NaCl. After incubation at 25 °C for 2 h, reactions were quenched with 500 µL ethyl acetate. After the ethyl acetate addition, reactions were mixed by vortex and centrifuged at 10,000 g for 5 min. Supernatants were then transferred into GC vials for reaction conversion and enantioselectivity analysis. For reactions testing the effect of pH, NaPi buffers with pH 4, 7, and 10 were used. For reactions investigating thermal stability, enzyme solutions at 10 mg/mL were first incubated at 50 °C for 30 min with all other reaction conditions remaining the same before cooling to 30°C to perform the reactions.

### Melting temperature

Thermal stability was assessed by intrinsic fluorescence using a NanoTemper Prometheus Panta, applying a 20–95 °C temperature ramp at 1 °C/min with samples at 0.2 mg/mL; Tm was determined from the inflection point (peak of the first derivative) of the 350/330nm unfolding transition

### Data curation for fine-tuning ESM-2

External deep mutational scanning datasets were used to benchmark representation and fine-tuning strategies before applying supervised modeling to Gre2. Three proteins were included: avGFP^34^, CreiLOV^33^, and Ube4b^35^. For each protein, curated variant sets were organized by number of mutations, including single, double, triple, quadruple, quintuple, and single-through-quintuple mutant training regimes. Datasets are described in detail in Supplementary Table 1 with train/validation splits described in Supplementary Table 2. Protein-specific held-out tests are described in Supplementary Table 3. Test sets were not used for model selection.

### Fine-tuning ESM-2

All benchmarks used the ESM-2 650M as the protein language model backbone. Three ESM-2 representations were compared: the final-layer CLS-token embedding, the mean-pooled final-layer embedding, and the full residue-level final-layer embedding. For residue-level embeddings, token embeddings were either flattened directly for full fine-tuning or projected through a learned bottleneck before flattening in the LoRA implementation. Further details are reported in Supplementary Table 4.

Three supervised modeling strategies were evaluated. First, frozen ESM-2 embeddings were used as inputs to a regularized linear regression model. This was the lowest-compute baseline because ESM-2 embeddings were precomputed and only the regression head was trained. Second, ESM-2 was fine-tuned with a multilayer perceptron regression head. In this implementation, the regression head and the final selected named ESM-2 parameter tensors were optimized jointly. Third, parameter-efficient LoRA fine-tuning was applied to ESM-2 attention modules, with adapters inserted into the query, key, value, and attention-output dense projections. The base model was frozen, and LoRA parameters were enabled only in the final 27 transformer blocks.

Model selection used validation mean-squared error with early stopping. Final performance was assessed on held-out test sets. Hyperparameters for each model family are reported in Supplementary Table 5. Scripts for finetuning ESM-2 are included in the Github repository.

### Data curation for VAE

We curated natural sequences related to Gre2 from the protein database^46^ with the Hidden-Markov model homology search tool Jackhmmer^47^ to obtain a multiple-sequence alignment (MSA) containing proteins related to Gre2 with a maximum of 2 iterations (N=2). We removed sequences less than 75% of the length of Gre2, removed sequences with an amino acid repeating 10 times in a row, and removed positions of the MSA not corresponding to Gre2. We reweighted the remaining 187,296 sequences with neighbors classified as having a Hamming distance/length of sequence greater than 0.98 to reduce phylogenetic bias from uneven sampling^17,48,49^. We withheld 100 sequences from the reweighted MSA to later assess VAE overfitting to the training set as a pseudo-test set given that these sequences are unlabeled. We randomly sampled the remaining 187,196 sequences with a 90/10 split for the training and validation sets.

### VAE pre-training

The VAE is trained by maximizing a modified version of the Evidence Lower Bound (ELBO) that effectively minimizes the Kullback–Leibler divergence between the variational approximation and the true posterior distribution^17^ as shown with equation (1):

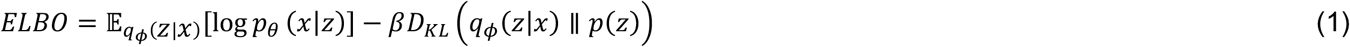

The first term can be considered a reconstruction loss that is computed using cross-entropy between an input one hot-encoded sequence and output likelihoods. β is the weight for the Kullback–Leibler divergence term *D*_*kl*_ with a prior distribution p(z) of N(0,I).

### Scoring sequence variants

For fine-tuned ESM-2 scoring, the raw model score was the predicted initial reaction rate for candidate sequence (x) with equation (1):

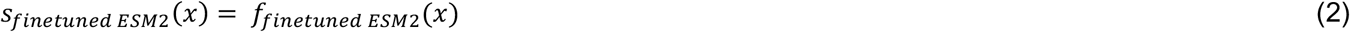

where (x) is a candidate sequence and *f*_*finetuned ESM2*_ (*x*) is the scalar prediction from the fine-tuned ESM-2 regression model.

To compute a VAE-based evolutionary likelihood for each sequence *x* = (*x*_1_, …, *x*_*L*_), we used the VAE decoder to compute the summed reconstruction cross-entropy across sequence positions. Because the VAE uses stochastic latent sampling during generation, we obtained deterministic scores by decoding from the mean of the approximate posterior, *z*_µ_, rather than from a sampled latent variable with equation (1):

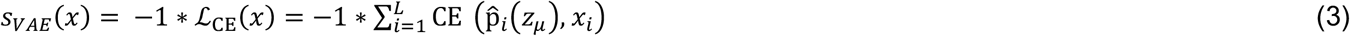

where 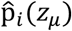 is the decoder-predicted categorical distribution at position *i* conditioned on *z*_µ_. Higher VAE scores correspond to lower reconstruction loss and greater evolutionary likelihood under the model.

SolubleMPNN was used to estimate whether each designed sequence was compatible with the Gre2 backbone while favoring soluble protein sequences. For each candidate sequence (x), we used SolubleMPNN in score-only mode conditioned on the Gre2 backbone structure, (B). Because SolubleMPNN scoring is stochastic, we scored each sequence (R) times and averaged the resulting backbone-conditioned sequence scores. The averaged score was multiplied by (-1), so that higher values corresponded to more favorable SolubleMPNN scores with equation (1):

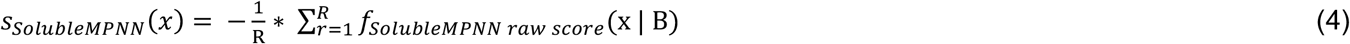

where *f*_*SolubleMPN raw score*_(x | B) is the raw SolubleMPNN score for sequence (x) conditioned on backbone (B) in scoring replicate (r). We used (R=5) scoring replicates per sequence.

### Multi-objective design of sequence variants

We generated co-optimized Gre2 sequence variants using simulated annealing adapted from previously described sequence-design workflows^14,30^. We designed variants with a fixed number of mutations while optimizing predicted reaction rate, evolutionary likelihood, and structural compatibility/solubility using fine-tuned ESM-2, a variational autoencoder (VAE), and SolubleMPNN^20^. All designs were generated in the context of the F85L parent sequence. Simulated annealing runs with 3 or 5 additional mutations corresponded to variants with 4 or 6 total mutations relative to the Gre2 sequence, respectively. The N-terminal methionine and F85L parent mutation were held fixed during optimization.

At each simulated annealing step, we proposed a new sequence by replacing a subset of mutations in the current sequence while maintaining the fixed total mutation count. The number of mutations replaced at each step was sampled from a Poisson distribution, with *λ*=1 when designing 4-mutation variants and *λ*=2 when designing 6-mutation variants. The number of replaced mutations was constrained to be at least one and less than the fixed total mutation count.

We first optimized each model objective separately to define objective-specific score ranges and a theoretical utopia point corresponding to the best value for each property. Fine-tuned ESM-2 was used to predict reaction rate, the VAE was used to estimate evolutionary likelihood, and SolubleMPNN was used to score sequence compatibility with the Gre2 structure.

For single-objective simulated annealing, the fitness function for candidate sequence (x) was the score from the model being optimized with equation (1).

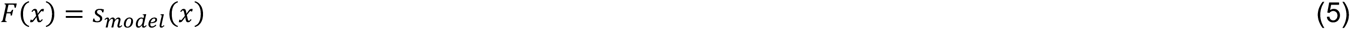

where *s*_*model*_(*x*) is the score from fine-tuned ESM-2, the VAE, or SolubleMPNN, using the sign convention that higher scores are more favorable. Proposed mutations that improved fitness were automatically accepted. All other proposed mutations were accepted according to the Metropolis criterion with equation (1):

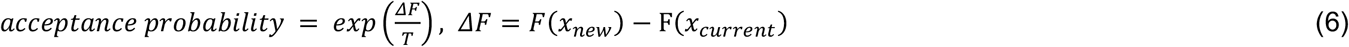

where *x*_*current*_ is the current sequence, *x*_*new*_ is the proposed sequence, and (T) is the simulated annealing temperature. The temperature decreased according to a logarithmic cooling schedule selected to encourage early exploration while increasingly favoring exploitation near the end of each trajectory.

We next optimized variants using fine-tuned ESM-2, the VAE, and SolubleMPNN simultaneously. Because the three model scores had different numerical scales, each score was min–max normalized before multi-objective combination with equation (1)

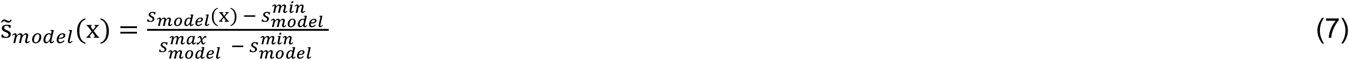

where *s*_*model*_(x) is the raw score from finetuned ESM-2, VAE, or SolubleMPNN, and 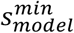 and 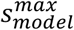 are the model-specific score bounds estimated from single-objective simulated annealing.

Multi-objective optimization was formulated as minimizing the Euclidean distance from the normalized utopia point, ((1,1,1)), corresponding to the best normalized value for predicted reaction rate, VAE evolutionary likelihood, and SolubleMPNN score with equation (1):

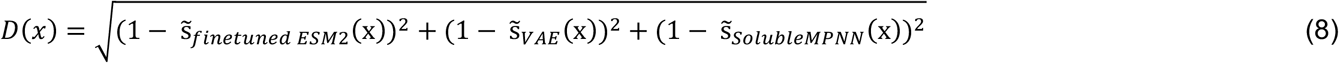

Variants closer to the utopia point had lower (D(x)) and therefore more favorable multi-objective scores. Proposed variants that reduced (D(x)) were automatically accepted. Variants that increased (D(x)) were accepted according to a distance-based Metropolis criterion with equation (1):

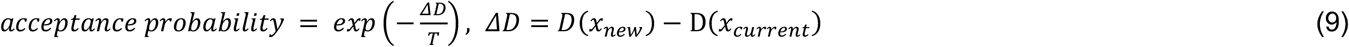

We selected sequence variants generated via simulated annealing simulations with all 3 model scores enhanced beyond Gre2 scores. This resulted in 1,219 sequence variants containing 4 or 6 mutations. Selected sequences were one-hot encoded by amino acid identity and clustered using the k-means++ algorithm. We then iteratively sampled sequences from each cluster to select diverse variants for experimental characterization (Figure 3g).

### Characterizing multi-site variant designs

For the small scale biocatalytic reactions, samples for gas-chromatography analysis were run on a HP 6890 GC with Supelco Beta DEX 225 (Part number 24348) 30 m × 0.25 mm × 0.25 µm column and FID detector. Program: 940°C equilibration time 1 min, 90°C hold time 5 min, ramp 50°6.8°C/min to 1490°C, 140°C hold time 4 minramp 1°C/min to 105°C, ramp °C/min 18°C/min 150°C, ramp 27.3°C/min to 40°C, 0.5 min150°C post run time 0.5 min. Retention times were: (S)-2-ethynyl-2-methylcyclohexan-1-one 14.75.32 min, (R)-2-ethynyl-2-methylcyclohexan-1-one 5.3915.0 min, (S)-2-ethynyl-2-methylcyclohexan-1-ol 15.25.48 min.

### Chemicals

Racemic, (R) or (S)-2-ethynyl-2-methylcyclohexan-1-one (CAS#: 65691-72-7) were previously reported^23^ and provided by Boehringer-Ingelheim. Acetophenone, (S)-1-phenylethan-1-ol, (R)-1-phenylethan-1-ol were purchased from Millipore Sigma, 4-phenylcyclohexanone was purchased from Fluorochem, cis-4-phenylcyclohexanol and trans-4-phenylcyclohexanol were purchased from Chempur. NADPH and NADH were purchased from Oriental Yeast Co., Ltd.

## Data Availability Statement

Datasets derived from previous deep mutational scanning studies^33–35^ and used for model benchmarking in this study can be downloaded from Hugging Face: https://huggingface.co/datasets/RomeroLab-Duke/protein-fitness-datasets-for-benchmarking-ft-esm2-strategies.

## Code Availability Statement

Code for training the variational autoencoder, finetuning ESM2 (650M), and performing multi-objective optimization of a protein via simulated annealing is available at the GitHub repository: https://github.com/RomeroLab/Multi-objective_enzyme_design under the Apache 2.0 license.

## Acknowledgements

We thank Michael Galant, Erich Spielvogel, and Leonhard Geist for producing and purifying the enzymes used in this study. We also thank Kristina Gueneva-Boucheva and Jon Reed for their contributions to and support of the foundational work that enabled the protein engineering presented here.

## Author Contributions

The following authors contributed to the ideation of the work presented here, HW, FB, JS, NP, LJK and PAR. Experimentation was done by authors NB, YS, JH, RL and XM. Manuscript drafting and editing was completed by authors NB, YS, SR, HW, LJK and PAR.

