## Supplementary Information for "Integrating complementary biological information for multi-objective enzyme engineering"

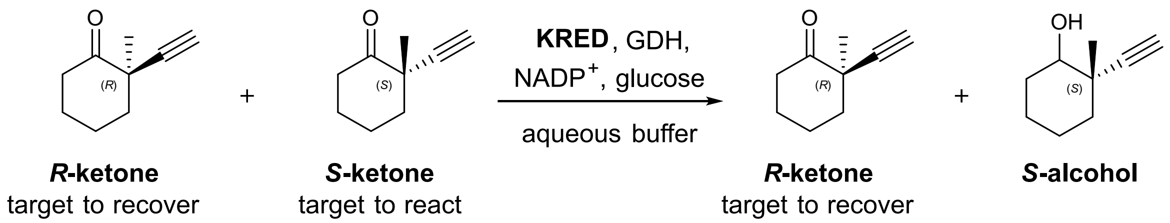


**Supplementary Figure 1:** Target kinetic resolution reaction

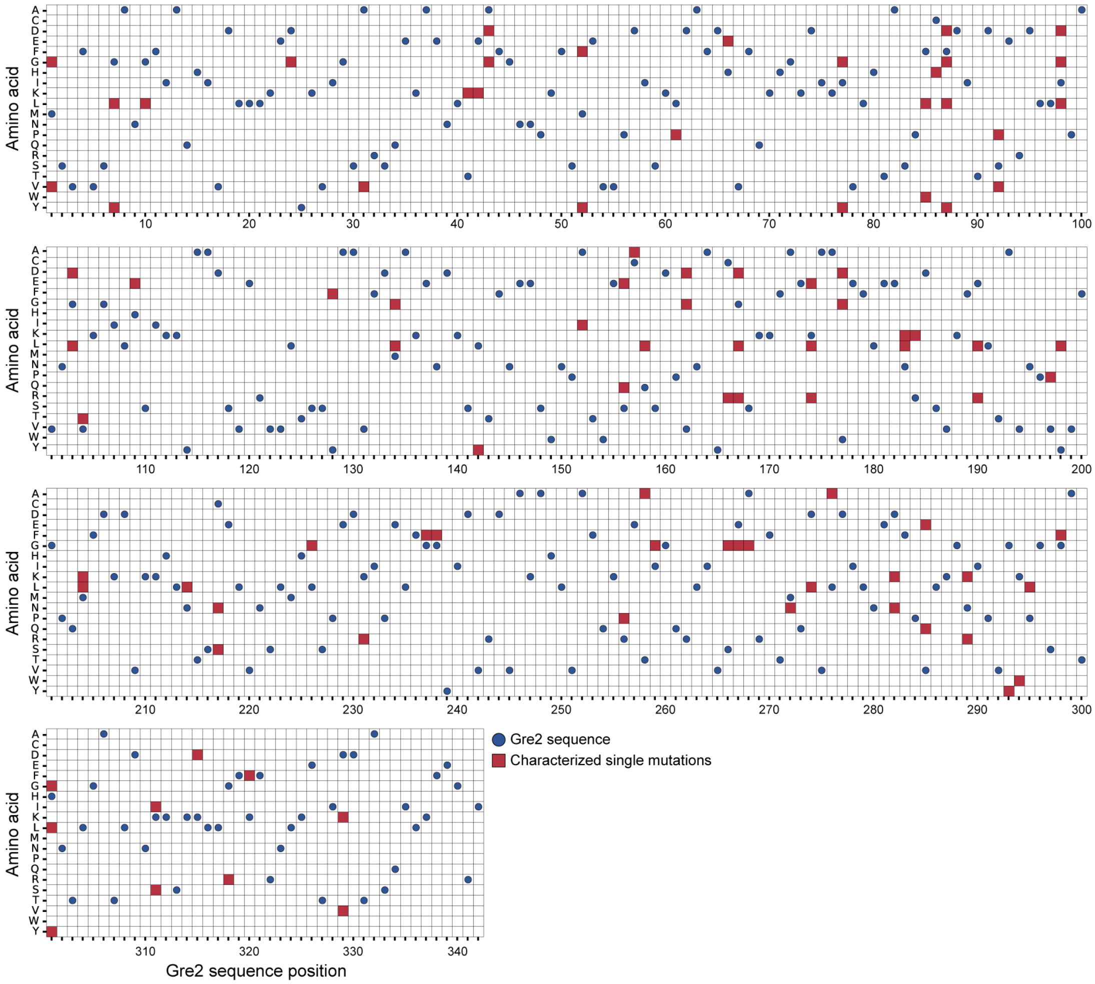


**Supplementary Figure 2:** Sparse sampling of the Gre2 single-mutant landscape. This heatmap with amino acid identity on the y-axis and Gre2 sequence position on the x-axis shows the initial Gre2 single-mutant dataset used for supervised activity modeling. Blue dots represent the Gre2 sequence. Red squares indicate a measured single amino-acid substitution. The dataset contained 99 characterized single mutants, covering 67 of 342 Gre2 positions and 99 of 6,498 possible single-amino-acid substitutions, corresponding to 19.59% positional coverage and 1.52% single-mutant coverage.

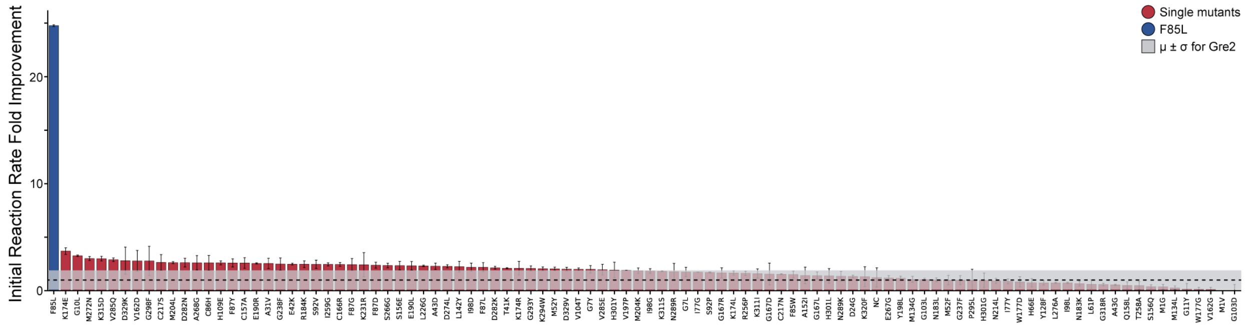


**Supplementary Figure 3:** Rank-ordered initial reaction-rate measurements for the 99 characterized Gre2 single mutants. Each bar represents one variant, with colors indicating qualitative expression category. The dashed horizontal line marks the wild-type Gre2 mean activity, and the shaded region indicates the relatively large standard deviation observed across wild-type measurements. Despite this WT variability, F85L was consistently among the strongest activity-enhancing variants and showed a clear improvement relative to WT Gre2. Most other single mutants showed activity near WT.

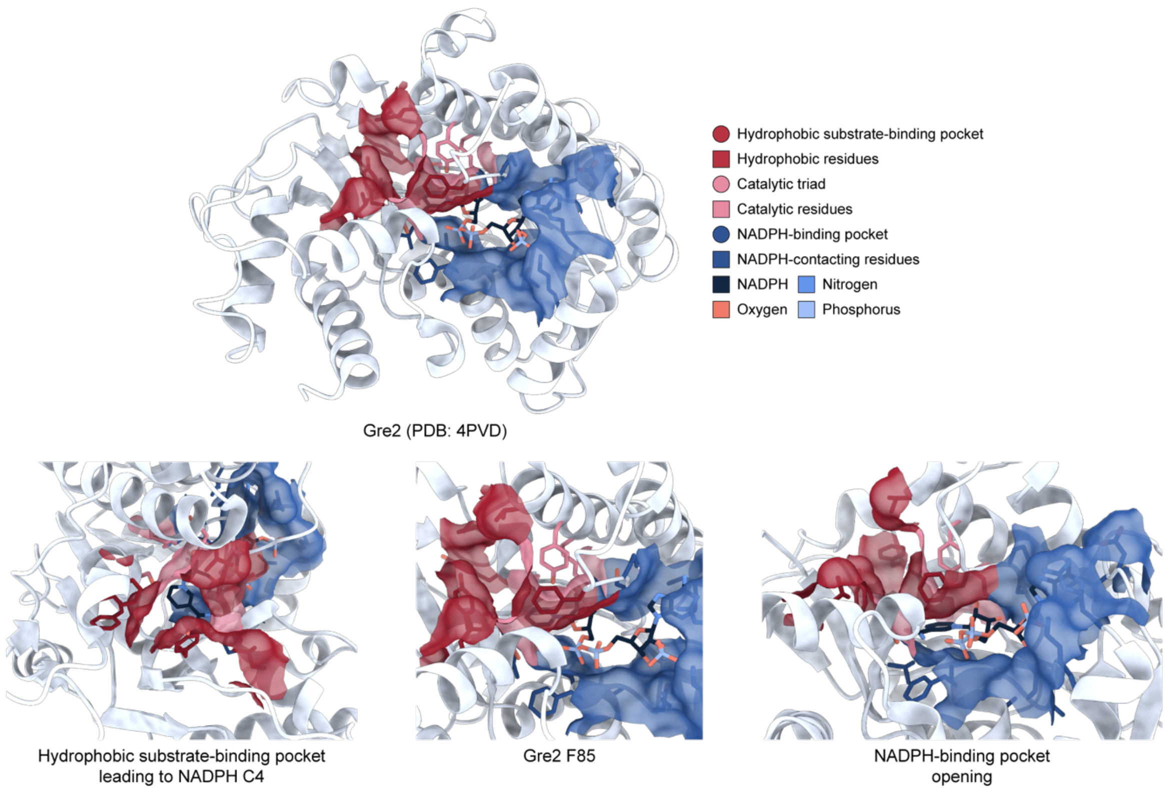


**Supplementary Figure 4:** Structural annotation of the Gre2 active site and location of F85. Annotated Gre2–NADPH structure highlighting the hydrophobic substrate-binding pocket (red) leading to the reaction center (NADPH C4) and NADPH-binding pocket (blue). F85 is located in the hydrophobic pocket near the substrate-binding channel and within the P84–C86 loop, providing structural context for why F85L could alter active-site geometry without globally disrupting the Gre2 fold. Multiple views are shown to illustrate the spatial relationship among NADPH C4, F85, and NADPH-binding pocket.

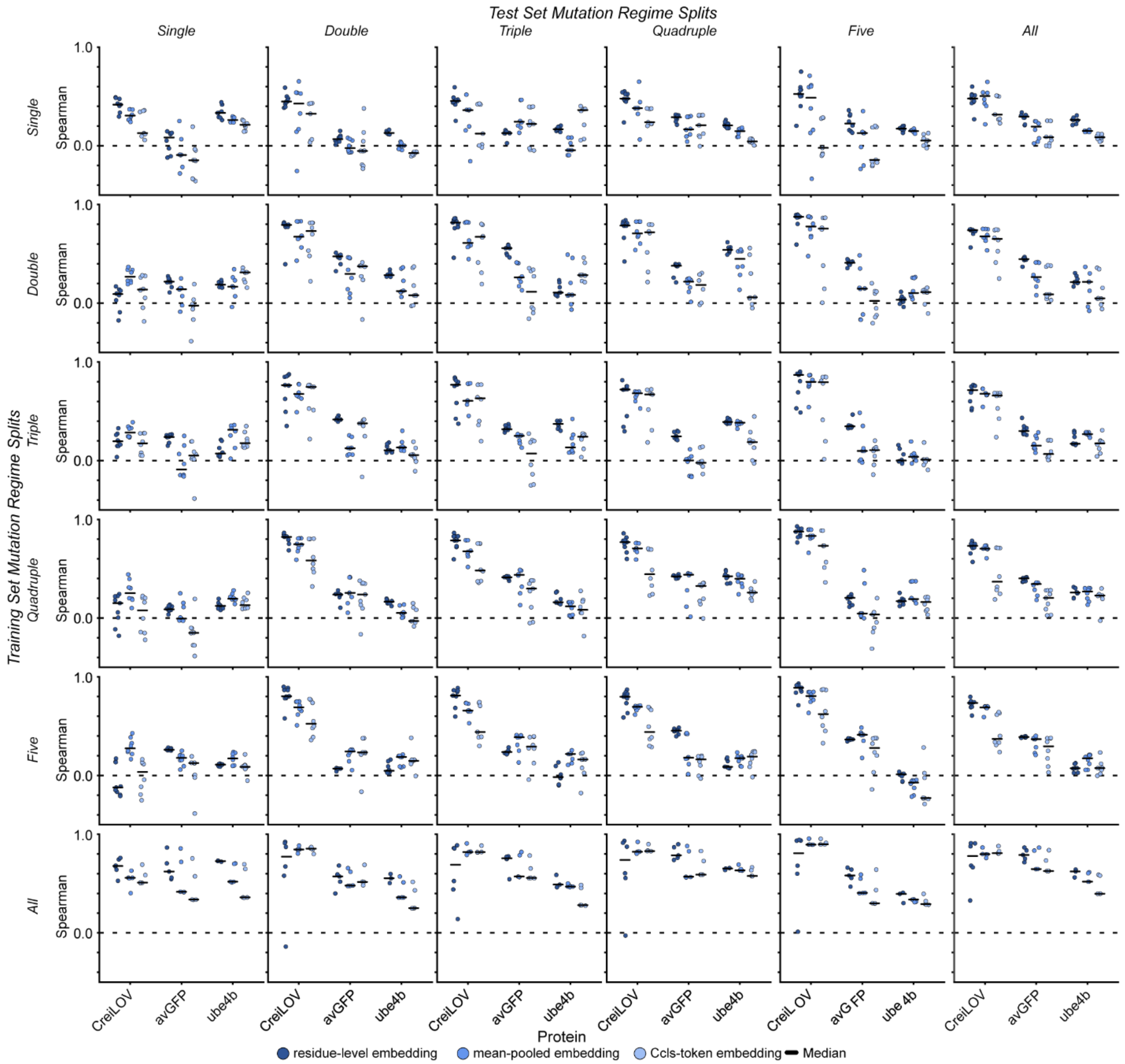


**Supplementary Figure 5:** ESM-2 representation strategies across external DMS datasets. Comparison of ESM-2 Cls-token, mean-pooled, and residue-level representations across CreiLOV, avGFP, and Ube4b low-data training regimes. Models were trained on mutation-regime-controlled datasets including single, double, triple, quadruple, quintuple, and single-through-quintuple mutant training sets as labeled on the rows. Each point represents model performance for a protein, mutation-order training regime, and random seed. Models were evaluated via Spearman rank correlations (y-axis) on held-out mutation-regime-controlled variants including single, double, triple, quadruple, quintuple, and single-through-quintuple mutant as labeled on the columns. Residue-level embeddings generally improved predictive performance in single-mutant training regimes (first row).

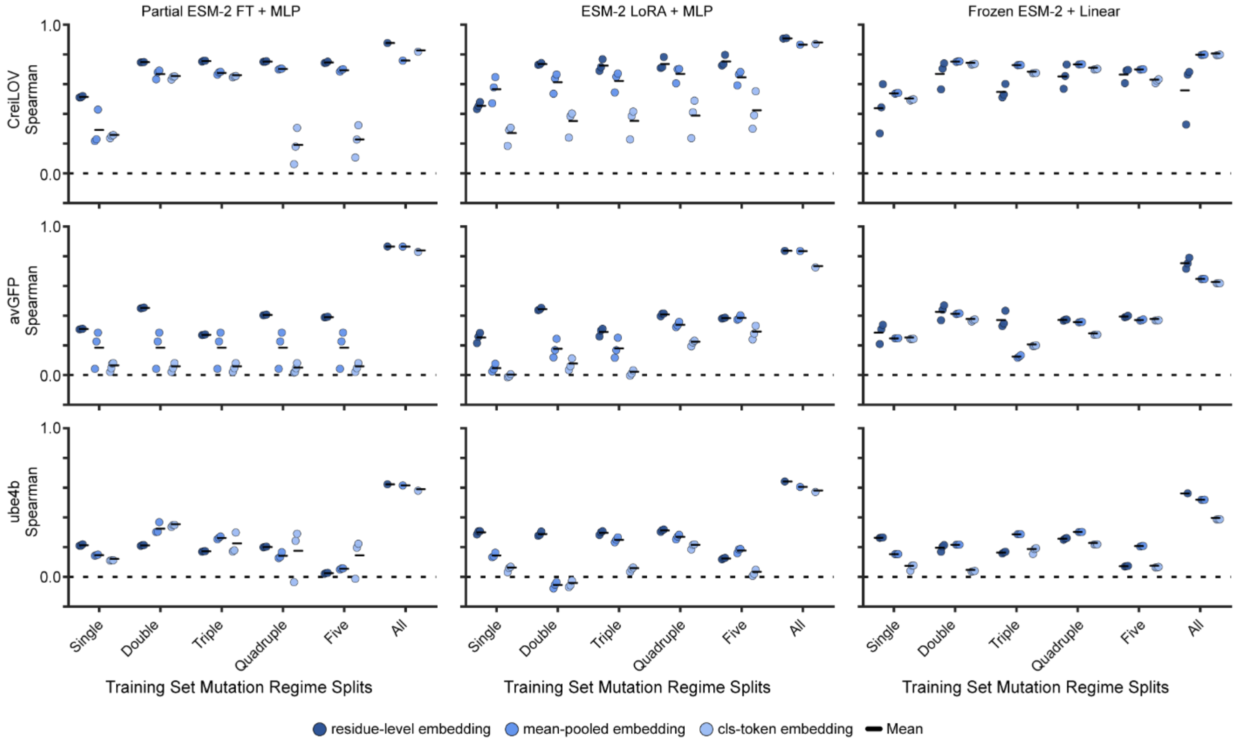


**Supplementary Figure 6:** ESM-2 fine-tuning strategies across external DMS datasets. Comparison of linear regression on frozen ESM-2 embeddings, LoRA-based parameter-efficient fine-tuning, and partial ESM-2 fine-tuning across external DMS benchmarks. Each point represents model performance for a protein, mutation-order training regime, and random seed. Parameter-efficient fine-tuning frequently improved held-out prediction relative to frozen-embedding linear models, supporting task-specific adaptation of ESM-2 in sparse sequence-function datasets.

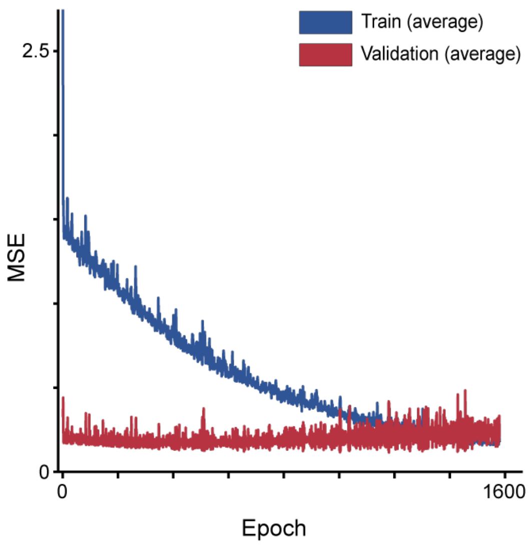


**Supplementary Figure 7:** Learning curve for fine-tuning ESM-2 on Gre2 activity measurements. Training and validation loss during supervised fine-tuning of ESM-2 on the 99 characterized Gre2 single mutants. The model was trained to predict experimentally measured initial reaction rate from sequence, with validation mean-squared error used for model selection and early stopping. The learning curve was used to monitor overfitting in the low-data Gre2 fine-tuning regime.

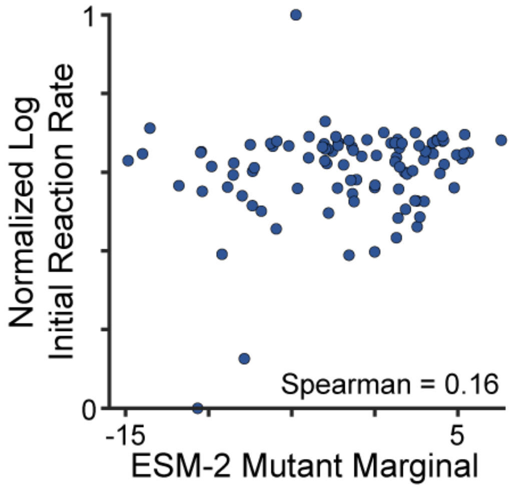


**Supplementary Figure 8:** Zero-shot ESM-2 baseline for Gre2 activity prediction. Relationship between zero-shot ESM-2 variant scores and experimentally measured initial reaction rates for the Gre2 single-mutant dataset. Zero-shot ESM-2 scores showed limited agreement with measured activity with a Spearman rank correlation of 0.16.

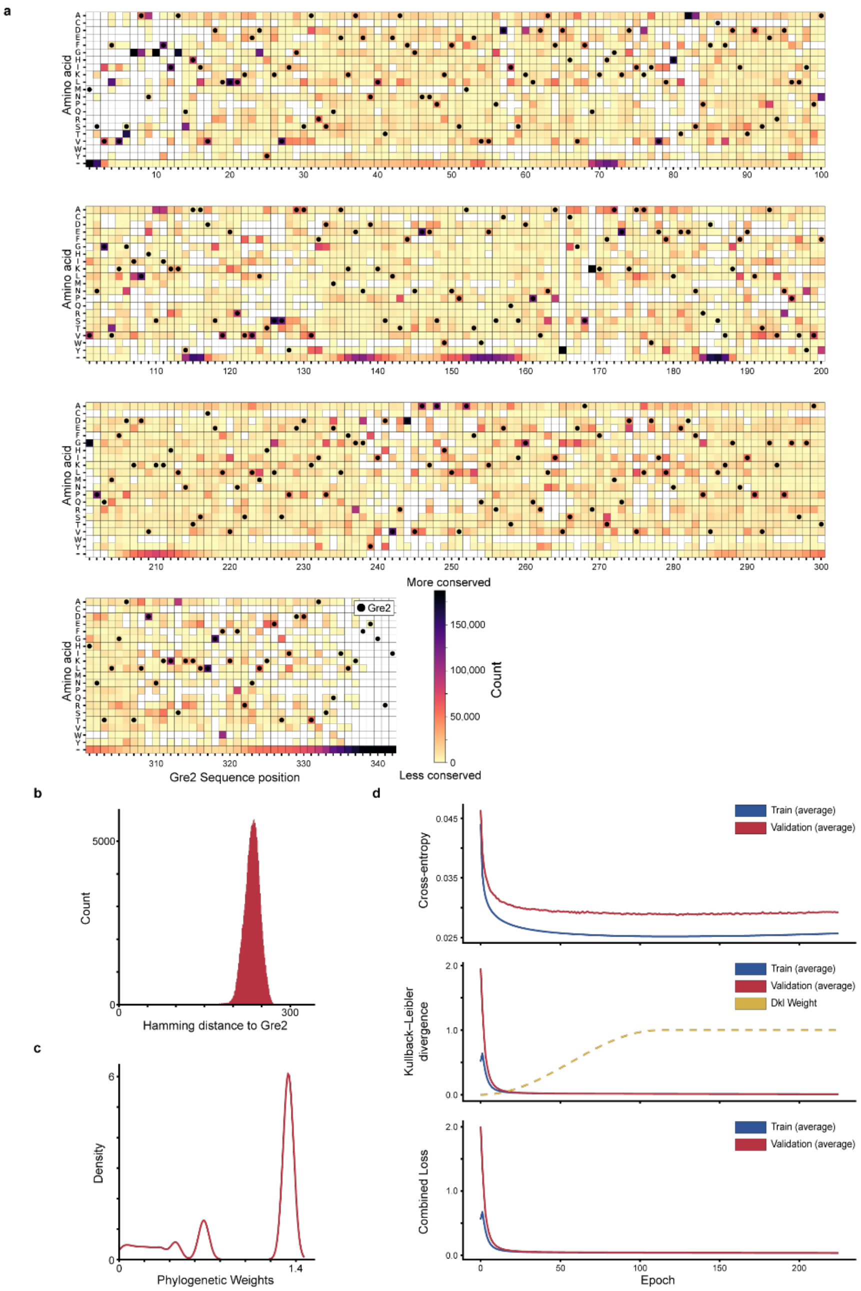


**Supplementary Figure 9:** VAE pre-training on natural Gre2 homologs. a, Amino-acid frequency heatmap for the curated multiple sequence alignment of natural Gre2-related homologs used to train the VAE. Amino-acid identity is shown on the y-axis and Gre2 sequence position on the x-axis, with color indicating the frequency of each amino acid at each aligned position. The Gre2 sequence is marked with black circles. b, Distribution of sequence identity for natural sequences in the curated MSA relative to the Gre2 sequence. c, Distribution of phylogenetic sequence weights used during VAE training to reduce bias from overrepresented homologous sequences. d, Training curves for VAE pre-training showing cross-entropy loss, Kullback-Leibler divergence, and combined VAE objective (ELBO).

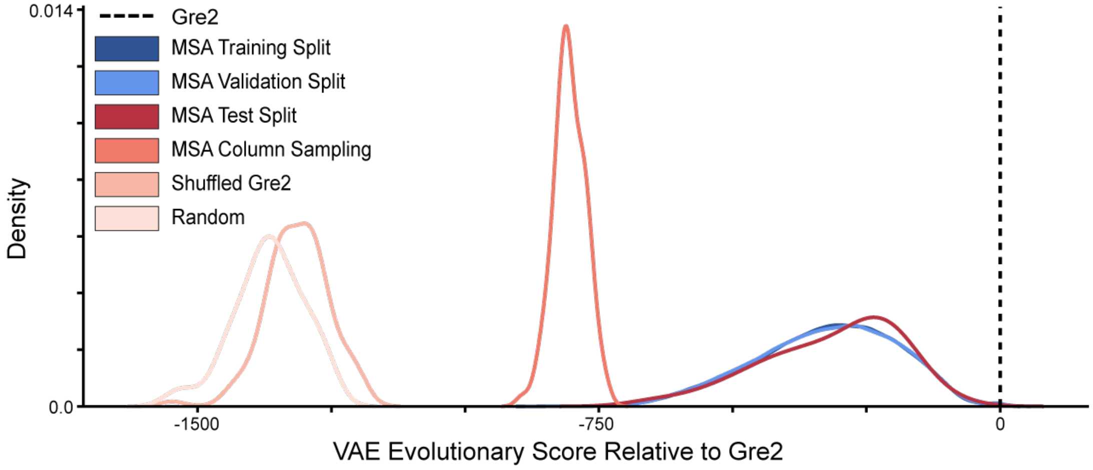


**Supplementary Figure 10:** VAE scores distinguish natural Gre2-family sequences from perturbed sequences. Distribution of VAE evolutionary scores across sequence classes used to assess model specificity. Natural Gre2-related sequences from the multiple sequence alignment (MSA) receive substantially higher scores than control sequence ensembles, including (i) column-resampled MSA sequences constructed by independently sampling residues from each alignment column to preserve site-wise frequencies while disrupting higher-order correlations, (ii) shuffled Gre2 sequences generated by randomly permuting amino acids while preserving amino-acid composition and gap structure, and (iii) fully random amino-acid sequences.

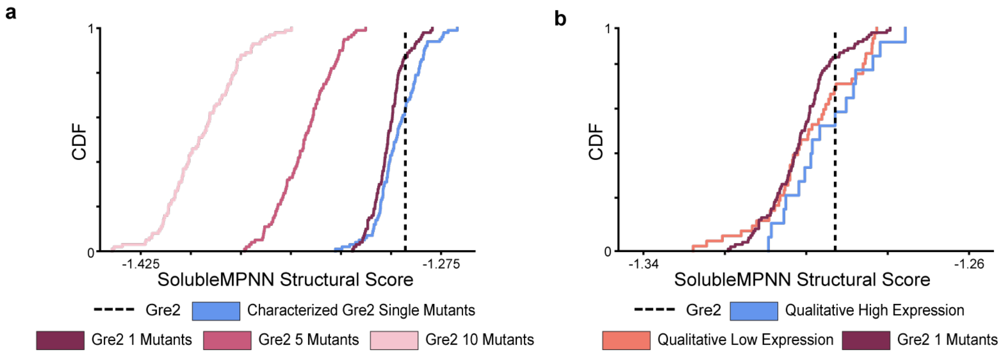


**Supplementary Figure 11:** SolubleMPNN provides a structure-based plausibility score for Gre2 variants. SolubleMPNN scores for Gre2 variants evaluated against the Gre2 structural scaffold. Random mutagenesis rapidly reduced structural plausibility, consistent with the model penalizing sequences that become less compatible with the folded structure. Previously characterized high-expression Gre2 single mutants showed partial separation from random single mutants, suggesting that SolubleMPNN captures structural or solubility-related information complementary to the activity-trained ESM-2 model.

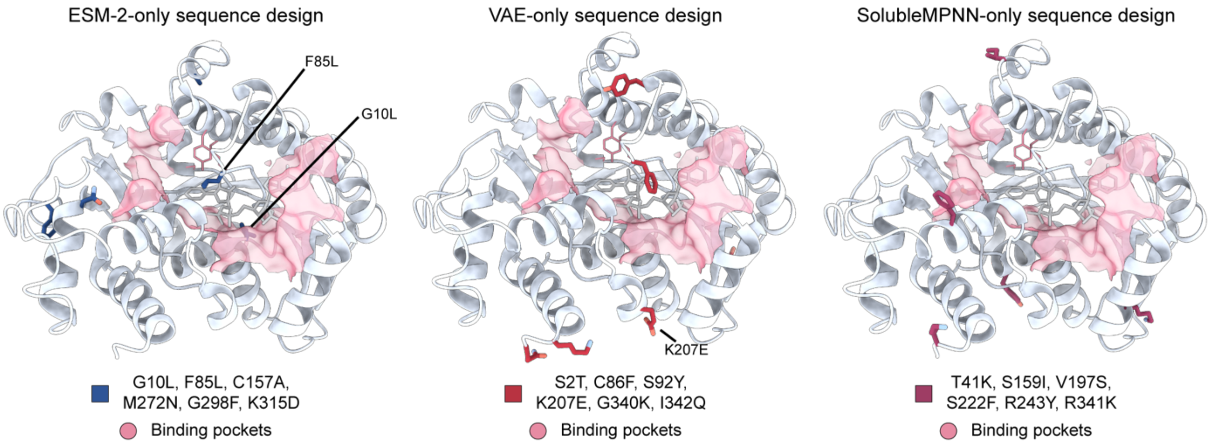


**Supplementary Figure 12:** Single-model sequence optimization produces structurally distinct design solutions. Structural mapping of mutations selected by single-objective simulated annealing using the fine-tuned ESM-2 activity model, the VAE evolutionary prior, or the SolubleMPNN structural prior. Structures were predicted via AlphaFold3[1]. The three independently optimized six-mutation designs contained no shared substitutions, indicating that the activity, evolutionary, and structural objectives prioritize distinct regions of Gre2 sequence and structure space.

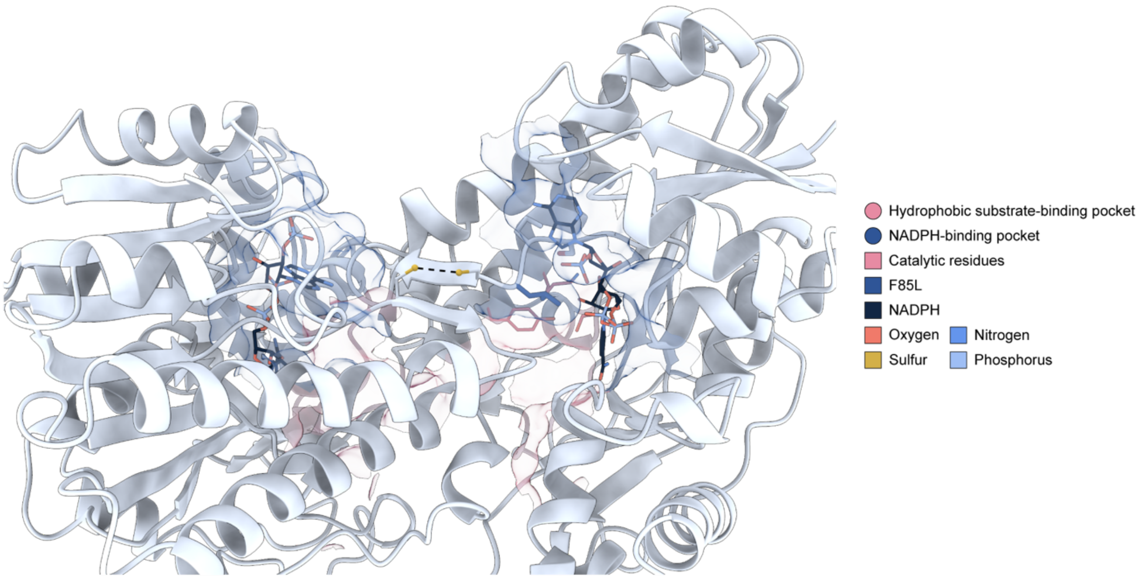


**Supplementary Figure 13:** Predicted Gre2 homodimer structure suggests a potential Cys86–Cys86 interchain disulfide. AlphaFold3[1]-predicted Gre2 homodimer structure highlighting the relative orientation of the two monomers, NADPH-binding pockets, substrate-binding regions, and catalytic residues. The Cys86 residues from opposing subunits are positioned near the dimer interface in a geometry consistent with a possible interchain disulfide bond, shown by the dashed line. This structural model suggests that Cys86 may contribute to dimer stabilization in Gre2.

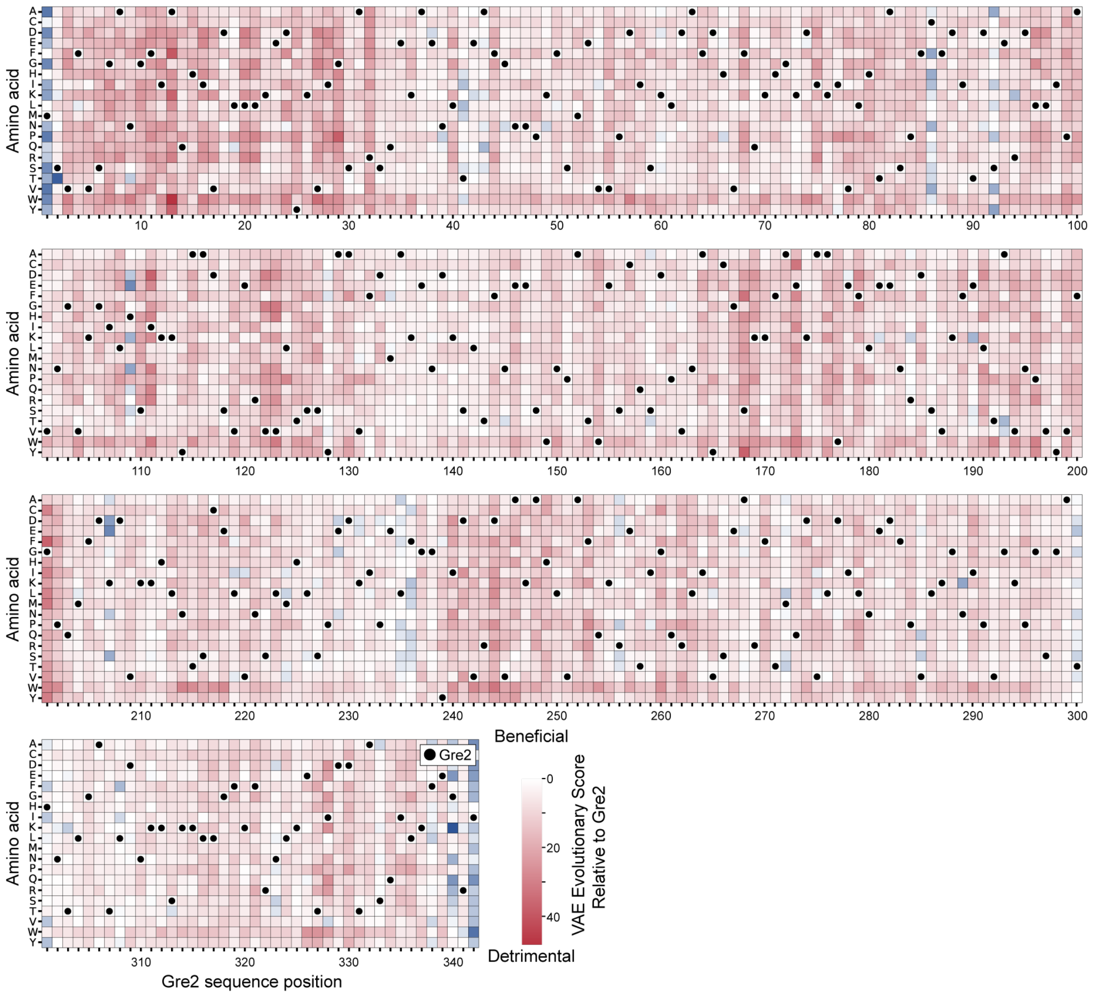


**Supplementary Figure 14:** VAE single-mutant preference landscape across Gre2. VAE-predicted evolutionary preferences for Gre2 single mutants across the full sequence. Each cell represents a possible amino-acid substitution at a Gre2 position, colored by the VAE score, with experimentally characterized or selected mutations marked. The landscape highlights mutations favored by the evolutionary prior, including H109E and K207E.

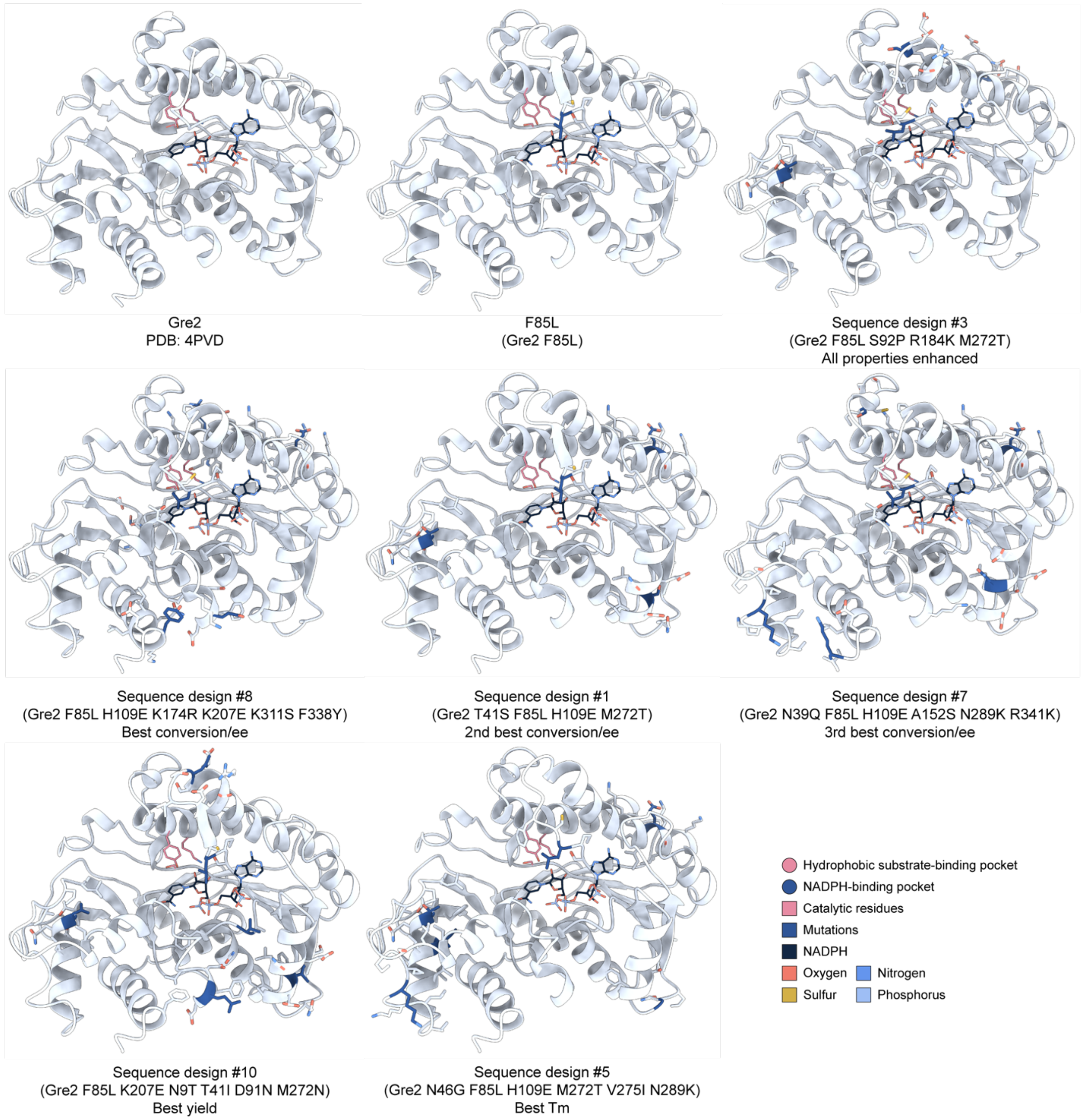


**Supplementary Figure 15:** Structural mapping of mutations in experimentally characterized designs. Gre2 crystal structure (PDB: 4PVD) and AlphaFold3[1]-predicted structural models of F85L and representative multi-objective sequence designs, including the all-properties-enhanced design #3, best reaction-performance design #8, second-best reaction-performance designs #1 and #7, best yield design #10, and best thermostability design #5. Predicted structures were aligned to the Gre2 crystal structure for comparison. High-performing variants all retain F85L but contain distinct additional mutations distributed across active-site-proximal, cofactor-proximal, substrate-proximal, and scaffold regions.

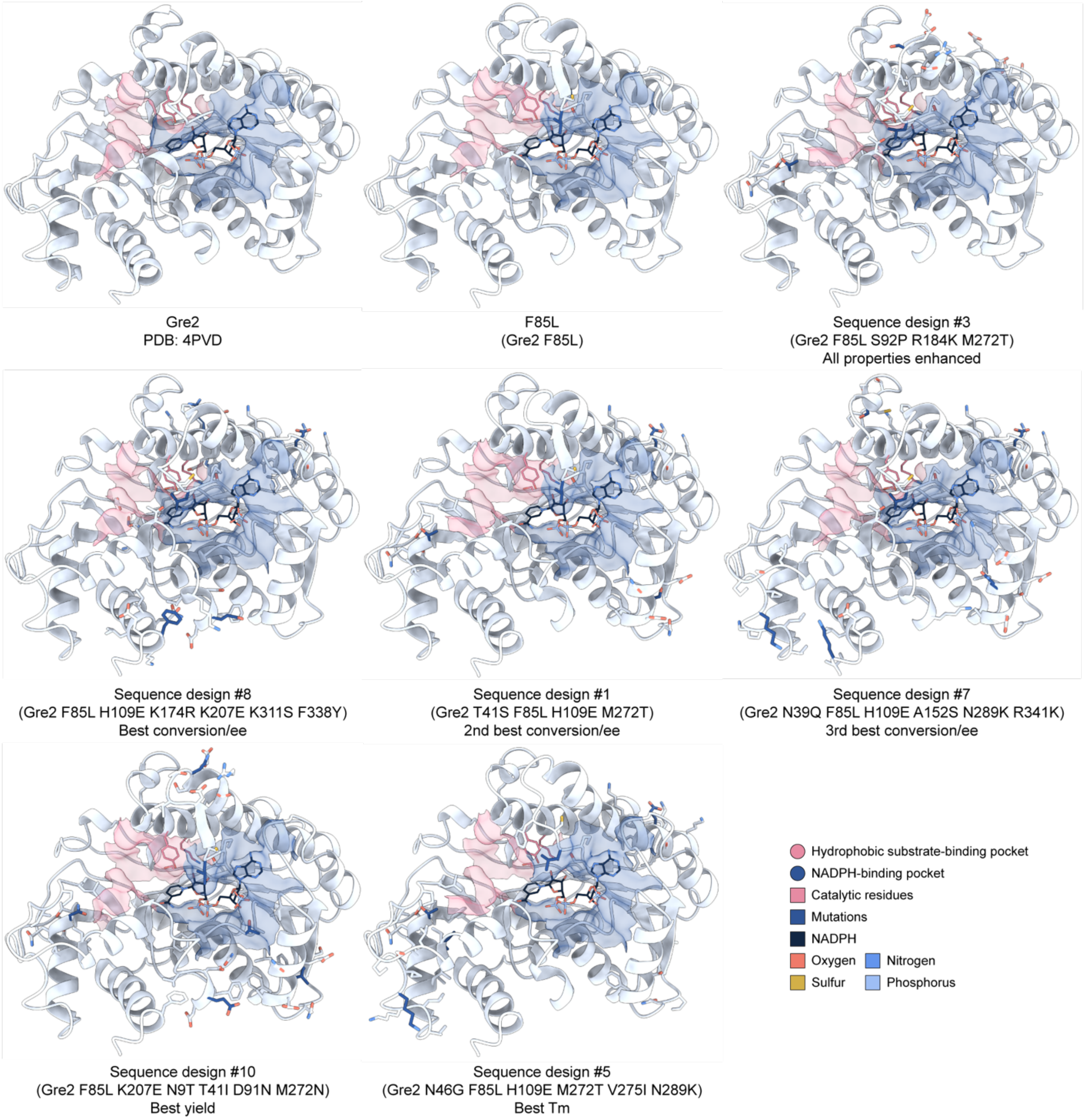


**Supplementary Figure 16:** Structural context of mutations relative to Gre2 active-site and cofactor-binding regions. Gre2 crystal structure (PDB: 4PVD) and AlphaFold3[1]-predicted structural models of F85L and representative multi-objective sequence designs, including the all-properties-enhanced design #3, best reaction-performance design #8, second-best reaction-performance designs #1 and #7, best yield design #10, and best thermostability design #5. Structures are annotated with the hydrophobic substrate-binding pocket, catalytic residues, NADPH-binding pocket, and NADPH to place design mutations in the context of functional regions involved in substrate recognition, cofactor binding, and catalysis. Predicted mutant structures were aligned to the Gre2 crystal structure for comparison.

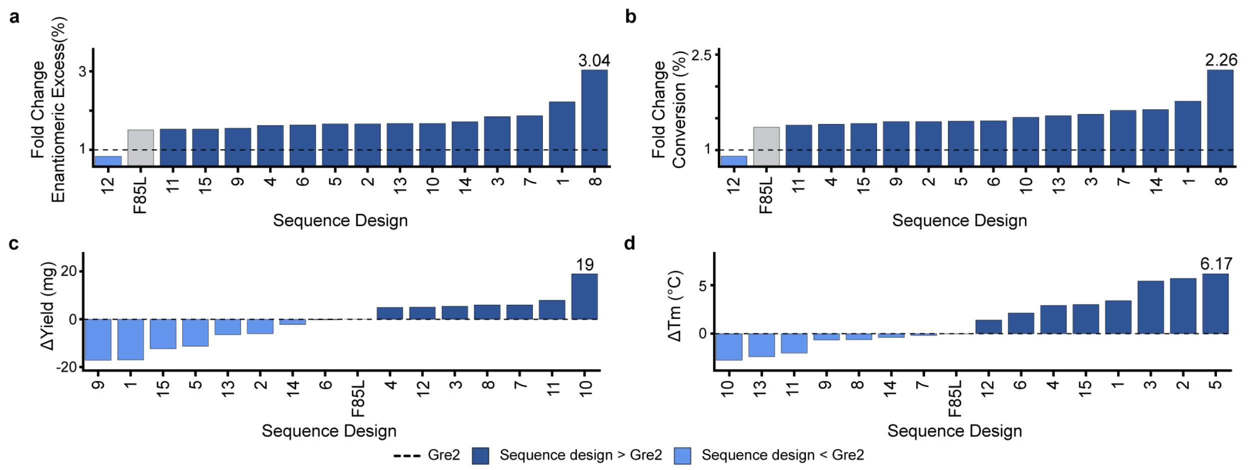


**Supplementary Figure 17:** Fold-change and absolute-property improvements for multi-objective Gre2 designs. Experimental improvements of multi-objective sequence designs relative to Gre2 and F85L across enantiomeric excess, conversion, protein yield, and thermal stability. Reaction-performance improvements are shown as fold changes, whereas yield and melting temperature are shown as absolute changes.

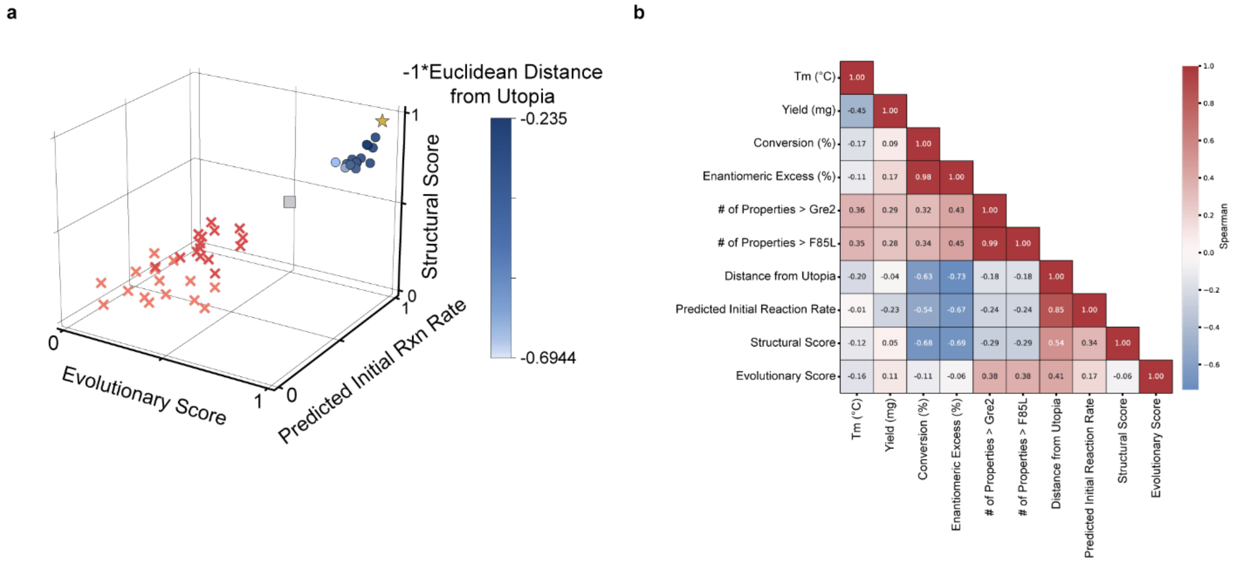


**Supplementary Figure 18:** Relationship between computational design scores and experimental outcomes. Comparison of computational multi-objective scores with experimentally measured improvements across the sequence design library. Candidate designs occupy an optimized region of predicted activity, evolutionary likelihood, and structural plausibility, but the number of experimentally improved properties did not map perfectly onto any single model score or Euclidean distance to the utopia point. Correlation analysis across model scores and experimental measurements indicates that no individual computational objective fully explains experimental performance, supporting diversity-aware sampling after multi-objective optimization.

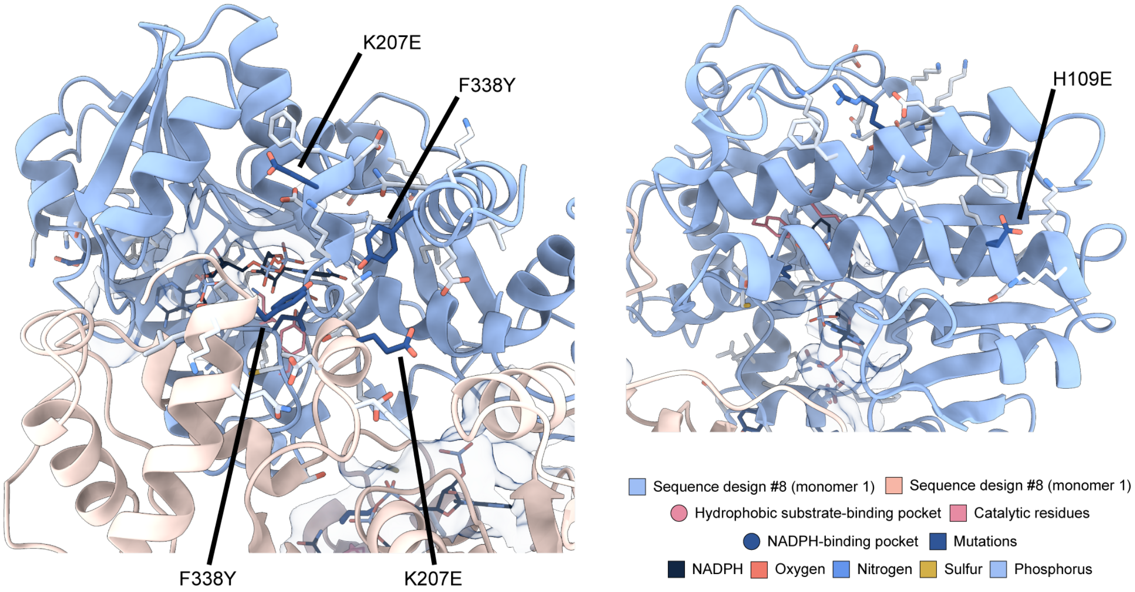


**Supplementary Figure 19:** Predicted homodimeric structural context of sequence design 8. AlphaFold3[1]-predicted homodimer structure of sequence design #8 shown in multiple structural snapshots. Design mutations are highlighted relative to NADPH, the NADPH-binding pocket, and the modeled dimer interface. K207E (and F338Y) are positioned near the dimer interface and the entrance to the NADPH-binding pocket. H109E lies on a helix surrounding the NADPH-binding pocket but is not near the dimer interface.

### **Supplementary Table 1:** CreiLOV, avGFP, and Ube4b DMS dataset composition

| Protein | Brief functional context | Size by number of mutations | | | | | Total |
|---|---|---|---|---|---|---|---|
| | | 1 mut. | 2 mut. | 3 mut. | 4 mut. | 5 mut. | 1–5 mut. |
| **CreiLOV** | LOV-domain fluorescent reporter; DMS readout measures cellular fluorescence as an engineered proxy for light-responsive function. | 2,206 | 176 | 978 | 3,565 | 9,603 | 16,528 |
| **avGFP** | Green fluorescent protein; DMS readout measures cellular fluorescence, integrating folding, maturation, and chromophore brightness. | 1,114 | 13,010 | 12,683 | 9,759 | 7,215 | 43,781 |
| **Ube4b** | Ubiquitination factor E4/E3 ligase domain; DMS readout reports ubiquitin-ligase activity or functional enrichment. | 932 | 54,507 | 31,748 | 8,782 | 1,903 | 97,872 |

Counts are sequence-variant counts for the 1–5 mutation regimes used in the idealized DMS training and evaluation setup.

### **Supplementary Table 2:** Training and validation splits

| Mut. regime | Training/validation pool | Train + val n | | | Train/val split | Model seeds |
|---|---|---|---|---|---|---|
| | | CreiLOV | avGFP | Ube4b | | |
| `1 Mut.` | WT + 100 single mutants | 101 | 101 | 101 | 90/10 | n = 3 |
| `2 Mut.` | WT + 100 double mutants | 101 | 101 | 101 | 90/10 | n = 3 |
| `3 Mut.` | WT + 100 triple mutants | 101 | 101 | 101 | 90/10 | n = 3 |
| `4 Mut.` | WT + 100 quadruple mutants | 101 | 101 | 101 | 90/10 | n = 3 |
| `5 Mut.` | WT + 100 five-mutant variants | 101 | 101 | 101 | 90/10 | n = 3 |
| `All` | WT + all remaining 1–5 mutation variants after held-out test removal | 16,279 | 43,532 | 97,623 | 90/10 | n = 1 |

Low-N subsets contain one WT sequence plus 100 sampled variants from the specified mutational regime. The `All` split used one model seed due to training cost.

### **Supplementary Table 3:** Fixed held-out test splits

| Mut. regime | Held-out test pool | Test n | | |
|---|---|---|---|---|
| | | CreiLOV | avGFP | Ube4b |
| `1 Mut.` | Single-mutant variants | 50 | 50 | 50 |
| `2 Mut.` | Double-mutant variants | 50 | 50 | 50 |
| `3 Mut.` | Triple-mutant variants | 50 | 50 | 50 |
| `4 Mut.` | Quadruple-mutant variants | 50 | 50 | 50 |
| `5 Mut.` | Five-mutant variants | 50 | 50 | 50 |
| `All` | **Total held-out 1–5 mutation variants** | **250** | **250** | **250** |

The same fixed held-out test set was used across the training/validation regimes for each protein.

### **Supplementary Table 4:** ESM-2 representations evaluated for sequence-function modeling.

| Representation | Implementation | Regression input size |
|---|---|---|
| CLS-token embedding | The first token embedding from the final ESM-2 hidden layer was used as a fixed-length sequence representation. | $d$ |
| Mean-pooled embedding | Final-layer token embeddings were averaged across the token dimension to produce a fixed-length sequence representation. | $d$ |
| Residue-level embedding | Final-layer token embeddings were retained for all positions. In the full fine-tuning model, these embeddings were flattened directly. In the LoRA model, token embeddings were first projected through a learned bottleneck and then flattened. | $(L+2)d$ or $(L+2)b$ |
| Precomputed CLS-token embedding | Frozen ESM-2 CLS-token embeddings were precomputed and used as input to a linear regression model. | $d$ |
| Precomputed mean-pooled embedding | Frozen ESM-2 mean-pooled embeddings were precomputed and used as input to a linear regression model. | $d$ |
| Precomputed Residue-level embedding | Frozen ESM-2 residue-level embeddings were precomputed and used as input to a linear regression model. | $(L+2)d$ |

$d$ denotes the ESM-2 hidden dimension (1280 for ESM-2 650M), $L$ denotes the protein sequence length, $L+2$ includes the start and end tokens used by the ESM-2 tokenizer, and $b$ denotes the learned bottleneck dimension used for residue-level LoRA models.

### **Supplementary Table 5:** Hyperparameters for ESM-2 650M fine-tuning.

| Hyperparameter | Frozen ESM-2 + linear regression | ESM-2 LoRA + MLP | Partial ESM-2 FT + MLP |
|---|---|---|---|
| ESM-2 trainable parameters | None | LoRA adapters enabled for final 27 of 33 transformer blocks | Final 27 named ESM-2 parameter tensors |
| Regression model | Single linear layer | Multilayer perceptron | Multilayer perceptron |
| Residue-level representation handling | Flattened embeddings | Per-token projection 640, followed by flattening | Flattened embeddings |
| LoRA target modules | N/A | Attention query, key, value, and output dense projections | N/A |
| LoRA rank, $r$ | N/A | 4 | N/A |
| LoRA $\alpha$ | N/A | 1 | N/A |
| LoRA dropout | N/A | 0.1 | N/A |
| Loss function | Mean-squared error | Mean-squared error | Mean-squared error |
| Optimizer | AdamW | AdamW | Adam |
| Learning rate | $1 \times 10^{-5}$ | $1 \times 10^{-6}$ for LoRA parameters; $1 \times 10^{-5}$ for MLP parameters | $1 \times 10^{-5}$ |
| Weight decay | $1 \times 10^{-5}$ | $1 \times 10^{-5}$ | $1 \times 10^{-5}$ |
| L1 penalty | $1 \times 10^{-3}$ | $1 \times 10^{-7}$ | N/A |
| Dropout | N/A | 0.1 | 0.1 |
| Gradient clipping | N/A | N/A | 3.0 |
| Scheduler | N/A | N/A | Cosine annealing warm restarts |
| EMA | N/A | N/A | Decay = 0.8 |
| Early stopping criterion | Validation MSE | Validation MSE | Validation MSE |
| Early stopping patience (epochs) | 500 if CreiLOV else 50 | 500 if CreiLOV else 100 | 500 if CreiLOV else 50 |
| Random seeds | 3, 7028, 88 | 3, 7028, 88 | 3, 7028, 88 |

The Partial ESM-2 FT implementation selected the final 27 named ESM-2 parameter tensors for optimization; this should not be interpreted as 27 complete transformer blocks. In contrast, the LoRA implementation explicitly enabled adapters in the final 27 transformer blocks.

**Supplementary Table 6:** Experimental characterization of 15 multi-objective Gre2 sequence designs for four process-relevant properties.

| Sequence design ID | Mutations | Tm [°C] | Yield [mg] | Conversion [%] | EE [%] |
|---|---|---|---|---|---|
| WT | — | 47.5 | 27.0 | 16.5 | 17.0 |
| F85L | F85L | 47.5 | 27.0 | 22.4 | 25.6 |
| 1 | T41S, F85L, H109E, M272T | 50.9 | 10.0 | 29.2 | 37.8 |
| 2 | N9T, F85L, S92P, M272T | 53.2 | 21.0 | 23.9 | 28.2 |
| 3 | F85L, S92P, R184K, M272T | 52.9 | 32.5 | 25.8 | 31.4 |
| 4 | F85L, H109E, K207E, M272T | 50.4 | 32.0 | 23.2 | 27.6 |
| 5 | N46G, F85L, H109E, M272T, V275I, N289K | 53.7 | 15.8 | 24.0 | 28.2 |
| 6 | N9T, E42K, F85L, K207E, M272T, K320S | 49.6 | 26.8 | 24.1 | 27.8 |
| 7 | N39Q, F85L, H109E, A152S, N289K, R341K | 47.3 | 33.0 | 26.8 | 31.8 |
| 8 | F85L, H109E, K174R, K207E, K311S, F338Y | 46.9 | 33.0 | 37.3 | 51.6 |
| 9 | F85L, H109E, R184K, V275I, N289K, D329A | 46.8 | 9.9 | 23.9 | 26.4 |
| 10 | N9T, T41I, F85L, D91N, K207E, M272N | 44.7 | 46.0 | 25.0 | 28.4 |
| 11 | T41L, F85L, H109E, F132Y, C166R, R256D | 45.5 | 35.0 | 23.0 | 26.0 |
| 12 | F85L, S92Y, H109E, C166H, K207E, D329T | 48.9 | 32.1 | 14.9 | 14.2 |
| 13 | F85L, H109E, R256E, A268E | 45.1 | 20.6 | 25.4 | 28.4 |
| 14 | K49N, F85L, H109E, E137Y, K207E, A268G | 47.1 | 24.8 | 27.0 | 29.2 |
| 15 | F85L, H109E, R184K, M272T | 50.5 | 14.7 | 23.4 | 26.0 |

Sequence design ID, amino acid substitutions, melting temperature (Tm), protein yield, conversion, and enantiomeric excess (ee) for Gre2 (WT), F85L, and the 15 multi-objective Gre2 sequence designs.

**Supplementary Table 7:** Performance of Gre2 sequence designs under pH challenge conditions.

| Sequence design ID | Mutations | pH | Conversion [%] | EE [%] |
|---|---|---|---|---|
| WT (Gre2) | — | 4.0 | 7.2 | 5.0 |
| WT (Gre2) | — | 7.0 | 16.5 | 17.0 |
| WT (Gre2) | — | 10.0 | 18.3 | 20.4 |
| 1 | T41S, F85L, H109E, M272T | 4.0 | 16.8 | 19.8 |
| 1 | T41S, F85L, H109E, M272T | 7.0 | 29.2 | 37.8 |
| 1 | T41S, F85L, H109E, M272T | 10.0 | 39.0 | 60.5 |
| 3 | F85L, S92P, R184K, M272T | 4.0 | 18.3 | 21.9 |
| 3 | F85L, S92P, R184K, M272T | 7.0 | 25.8 | 31.4 |
| 3 | F85L, S92P, R184K, M272T | 10.0 | 37.8 | 58.0 |
| 7 | N39Q, F85L, H109E, A152S, N289K, R341K | 4.0 | 20.3 | 25.0 |
| 7 | N39Q, F85L, H109E, A152S, N289K, R341K | 7.0 | 26.8 | 31.8 |
| 7 | N39Q, F85L, H109E, A152S, N289K, R341K | 10.0 | 42.9 | 71.6 |
| 8 | F85L, H109E, K174R, K207E, K311S, F338Y | 4.0 | 10.7 | 9.0 |
| 8 | F85L, H109E, K174R, K207E, K311S, F338Y | 7.0 | 37.3 | 51.6 |
| 8 | F85L, H109E, K174R, K207E, K311S, F338Y | 10.0 | 26.8 | 33.0 |
| 17 | M1G, F85L, K174E | 4.0 | 17.5 | 20.9 |
| 17 | M1G, F85L, K174E | 7.0 | 28.4 | 35.8 |
| 17 | M1G, F85L, K174E | 10.0 | 39.3 | 61.1 |

Conversion and enantiomeric excess (ee) measured for wild-type Gre2 and selected sequence designs across the evaluated pH conditions.

**Supplementary Table 8:** Performance of Gre2 sequence designs under temperature challenge conditions.

| Sequence design ID | Mutations | Temperature [°C] | Conversion [%] | EE [%] |
|---|---|---|---|---|
| WT (Gre2) | — | 30.0 | 16.5 | 17.0 |
| WT (Gre2) | — | 50.0 | 11.5 | 10.4 |
| 1 | T41S, F85L, H109E, M272T | 30.0 | 29.2 | 37.8 |
| 1 | T41S, F85L, H109E, M272T | 50.0 | 1.3 | 1.3 |
| 3 | F85L, S92P, R184K, M272T | 30.0 | 25.8 | 31.4 |
| 3 | F85L, S92P, R184K, M272T | 50.0 | 13.1 | 15.0 |
| 7 | N39Q, F85L, H109E, A152S, N289K, R341K | 30.0 | 26.8 | 31.8 |
| 7 | N39Q, F85L, H109E, A152S, N289K, R341K | 50.0 | 0.0 | 0.0 |
| 8 | F85L, H109E, K174R, K207E, K311S, F338Y | 30.0 | 37.3 | 51.6 |
| 8 | F85L, H109E, K174R, K207E, K311S, F338Y | 50.0 | 12.4 | 10.4 |
| 17 | M1G, F85L, K174E | 30.0 | 28.4 | 35.8 |
| 17 | M1G, F85L, K174E | 50.0 | 0.9 | 0.9 |

Conversion and enantiomeric excess (ee) measured for wild-type Gre2 and selected sequence designs across the evaluated reaction temperatures.
